# Structure of the human Duffy antigen receptor

**DOI:** 10.1101/2023.07.09.548245

**Authors:** Shirsha Saha, Basavraj Khanppnavar, Jagannath Maharana, Heeryung Kim, Carlo Marion C. Carino, Carole Daly, Shane Houston, Poonam Kumari, Prem N. Yadav, Bianca Plouffe, Asuka Inoue, Ka Young Chung, Ramanuj Banerjee, Volodymyr M. Korkhov, Arun K. Shukla

## Abstract

The Duffy antigen receptor, also known as FY glycoprotein or CD234, is a seven transmembrane protein expressed primarily at the surface of red blood cells, which displays promiscuous binding to multiple chemokines. Not only does it serve as the basis of the Duffy blood group system but it also acts as the primary attachment site for malarial parasite *Plasmodium vivax* on erythrocytes and as one of the nucleating receptors for the pore forming toxins secreted by *Staphylococcus aureus*. Despite a predicted 7TM architecture and efficient binding to a spectrum of chemokines, it fails to exhibit canonical second messenger response such as calcium release, likely due to a lack of G protein coupling. Unlike prototypical GPCRs and β-arrestin-biased atypical chemokine receptors, the Duffy antigen receptor also appears to lack β-arrestin binding, making it an enigmatic 7TM chemokine receptor. In order to decipher the molecular mechanism of this intriguing functional divergence exhibited by the Duffy antigen receptor, we have determined its cryo-EM structure in complex with a C-C type chemokine, CCL7. The structure reveals a relatively superficial binding mode of CCL7, with the N-terminus of the receptor serving as the key interaction interface, and a partially formed orthosteric binding pocket lacking the second site for chemokine recognition compared to prototypical chemokine receptors. The structural framework allows us to employ HDX-MS approach to uncover ligand-induced structural changes in the receptor and draw important insights into the promiscuous nature of chemokine binding. Interestingly, we also observe a dramatic shortening of TM5 and 6 on the intracellular side, compared to prototypical GPCRs, which precludes the coupling of canonical signal-transducers namely G proteins, GRKs and β-arrestins, as demonstrated through extensive cellular assays. Taken together, our study uncovers a previously unknown structural mechanism that imparts unique functional divergence on the 7TM fold encoded in the Duffy antigen receptor while maintaining its scavenging function and should facilitate the designing of novel therapeutics targeting this receptor.

## Main

The Duffy antigen, originally referred to as Fy glycoprotein, was first identified as a blood antigen expressed on the surface of erythrocytes^1, 2^ (Fig.1a), and the polymorphism displayed by this gene was used to classify the Duffy blood group system^2^. Consequently, it was found to be expressed by additional cell types such as epithelial cells of lung and kidney^3^, endothelial cells of capillaries^4^, hair cells of cochlea^5^, airway smooth muscle cells^6^, and selected regions of brain^7–9^. Based on a predicted seven transmembrane (7TM) topology and its ability to recognize multiple chemokines (Fig.1b), it was subsequently referred to as the Duffy Antigen Receptor for Chemokines (DARC), and categorized as a chemokine receptor in the superfamily of G protein-coupled receptors (GPCRs)^10, 11^. Strikingly however, it does not exhibit productive G protein-coupling as measured using second messenger response with calcium release as a readout, making it an enigmatic seven transmembrane receptor (7TMR)^9–12^.

DARC expressed on the surface of erythrocytes serves as the primary receptor for the malarial parasites *Plasmodium vivax* and *Plasmodium knowlesi*^13^, and these pathogens latch on to erythrocytes via the interaction of their Duffy Binding Protein (DBP) with DARC^14–17^ (Fig. 1c). Previous studies have demonstrated that DBP-DARC interaction is mediated primarily by the N-terminus of DARC, while the transmembrane core appears to be dispensable for the same^14–18^. Interestingly, a sub-set of population of sub-Saharan descent lack DARC expression on their erythrocytes, and this confers them resistance to *P. vivax* infection^19^. Furthermore, several of the bicomponent pore-forming toxins (PFTs), such as leukotoxin ED (LukED) and γ-haemolysin AB (HlgAB) secreted by *Staphylococcus aureus* also interact with DARC on erythrocytes, primarily through its N-terminus^20, 21^ (Fig. 1d). This interaction presumably serves as the anchor and nucleation site facilitating the oligomerization of the bicomponent PFTs, leading to subsequent pore formation, cell lysis and release of hemoglobin. Emerging data now show that PFTs can also lead to cell lysis and tissue damage by targeting DARC expressed on endothelial cells^22^. A recent study has suggested that some PFTs may also compete with chemokine binding to DARC, and induce structural changes in the receptor upon binding^23^. Moreover, several studies have also demonstrated an intriguing link between DARC expression and HIV susceptibility, where the absence of DARC appears to slow down the disease progression, although some studies have also suggested a direct role of DARC in promoting trans-infection of effector cells by HIV-1 virions^24–27^. These indications make DARC an important therapeutic target from multiple perspectives including malarial infection, anti-microbial resistance, and HIV infection.

**Fig. 1.**
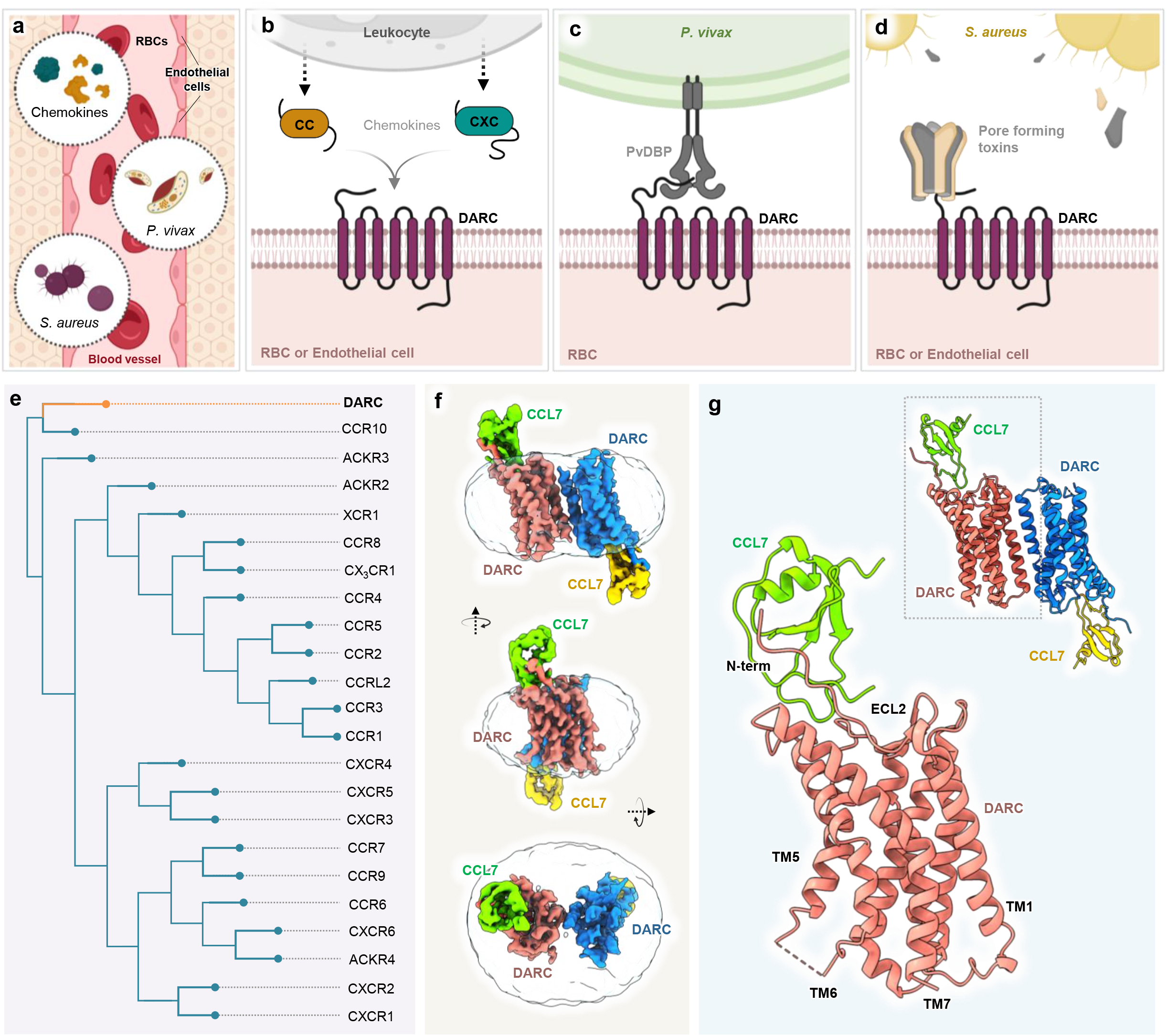
Structure of the human Duffy antigen receptor. **a.** DARC is expressed in erythrocytes and endothelial cells, and provides the docking site for various chemokines and pathogens. **b.** CC and CXC chemokines released from leukocytes reportedly bind to DARC. **c.** The malarial parasite *Plasmodium vivax* interacts and evades RBCs through the interaction of PvDBP and DARC. **d.** Multiple pore-forming toxins secreted by the pathogen *Staphylococcus aureus* also target DARC receptor for pore formation. **e.** Phylogenetic analysis of all chemokine receptors. **f, g.** Cryo-EM structure of CCL7 bound DARC dimeric complex shown in surface (f) and as cartoon representation (g) (Salmon and blue, DARC protomers; green and yellow, CCL7). Cryo-EM maps have been shown in different orientations, and a model of the monomeric complex of CCL7-DARC is highlighted as zoomed-in view.

While chemokine receptors often display ligand promiscuity in terms of recognition and activation by multiple chemokines, DARC represents the most striking example of promiscuous chemokine binding with its ability to recognize both C-C and C-X-C type chemokines, the only 7TMR capable of doing so^28–31^ (Fig. 1b). It has been proposed that such promiscuous binding to chemokines allows it to scavenge the ligands and establish a chemokine gradient that may have functional implications^32^. A recent study has also demonstrated that DARC expressed on the surface of developing erythrocytes robustly binds CXCL12 but the same receptor when present on mature erythrocytes loses the ability to recognize CXCL12, suggesting a complex regulatory mechanism driving ligand recognition, possibly through post-translational modifications and/or conformational changes^33^. In addition, a dimeric form of CXCL12 appears to have stronger binding affinity to DARC compared to the monomeric form^34^. DARC is also characterized as a member of the so-called atypical chemokine receptor sub-group (ACKRs), which lack functional G protein-coupling despite exhibiting a conserved 7TM architecture like GPCRs and robust chemokine binding, and is referred to as ACKR1^10, 35–38^. Interestingly however, while most of the ACKRs such as ACKR2, ACKR3 and ACKR4 exhibit robust β-arrestin-coupling upon agonist-stimulation^38, 39^, the same remains primarily unexplored for DARC. Despite the important contributions of DARC in regulating chemokine homeostasis and pathogenic infections, and a distinct functional manifestation compared to prototypical GPCRs and chemokine receptors, the structural characterization of this receptor remains primarily unexplored and represents an important knowledge gap.

Here, we present a cryo-EM structure of DARC in complex with CCL7, which reveals a distinct mode of chemokine interaction with the receptor compared to prototypical chemokine receptors in terms of second binding site, and provides a molecular mechanism to rationalize the lack of canonical transducer-coupling at this receptor while potentially maintaining its ligand scavenging function. This study elucidates a fundamental mechanism encoding functional divergence in 7TM proteins, and provides a previously missing framework paving the way for designing novel therapeutics.

## Overall structure of CCL7-DARC

Phylogenetic analysis positions DARC with prototypical chemokine receptor CCR10, with ACKR2 and ACKR3 being the next closest neighbors when compared to chemokine binding GPCRs (Fig. 1e) and GPR182 and GPR82 being the closest neighbors in addition to ACKR3 when compared to the entire GPCRome (Extended Data Fig. 1). As mentioned earlier, ACKR2 and ACKR3 are also chemokine receptors, however, they exhibit selective coupling to β-arrestins without any detectable G protein activation^40, 41^. We expressed and purified full length, wild-type DARC in the presence of a C-C type chemokine, CCL7, and a nanobody (Nb52) targeted against the N-terminus of DARC^42^. The purified receptor exhibited two distinct populations on size exclusion chromatography, which presumably represent monomeric and dimeric states of the ligand-receptor complexes (Extended Data Fig. 2a-b). Considering the small size of CCL7-DARC complex, we subjected the dimeric population to cryo-EM and observed an overall monodisperse population with 2D class averages showing a non-physiological arrangement of the ligand-receptor complexes (Extended Data Fig. 2c-d). Subsequently, we determined the structure of CCL7-DARC at an estimated resolution of 3.8Å, with clearly discernible densities for the receptor and ligand (Fig. 1f-g, Extended Data Fig. 2e-h, Extended Data Fig. 3a-b, and Extended Data Table 1). However, we did not obtain any densities for Nb52. The two protomers show a typical GPCR fold with a 7TM architecture, and have a nearly identical structure with an RMSD of <0.5Å (Fig. 1f-g).

## A distinct mode of ligand binding in DARC

We observe that CCL7 makes extensive contacts with the N-terminus of the receptor with some contribution from the extracellular side of TM6, TM7, and 3^rd^ extracellular loop (ECL3) (Fig. 2a-c and Extended Data Table 2). The N-terminal residues from Glu46 to Asn55 of DARC are positioned in a groove formed by CCL7, and this interaction is stabilized by several hydrogen bonds and other non-bonded contacts (Fig. 2a-b and Extended Data Table 2). Specifically, the N-terminal residues of DARC from Glu46 to Leu57 form an extended loop, which orients itself onto a docking site on CCL7 formed by the N-terminal loop residues Ser^8^ to Lys^18^ and the β-strand3 residues Lys^49^ to Ala^53^ (Fig. 2b). It is also interesting to note that the disulfide bridge in DARC between Cys51 in the N-terminus and Cys276 in TM7 is packed against a disulfide bridge formed between Cys^12^ and Cys^52^ of CCL7, and this helps the N-terminus of DARC align in the CCL7 groove (Fig. 2c). Chemokine receptors typically engage chemokines through two binding sites known as chemokine recognition site 1 and 2 (CRS1 and 2)^43^. However, CCL7 interaction with DARC appears to engage only CRS1 without any significant interaction with CRS2 (Fig. 2a) This is evident upon comparing CCL7-DARC interface with a previously determined CCL2-CCR2 complex through a cross-sectional area, where the binding pocket corresponding to CRS2 is empty (Fig. 2d). This is further apparent upon comparison with other chemokine-chemokine receptor complexes in G protein-bound, active conformation (Fig. 2e). Interestingly, the lack of CRS2 binding site observed here is reminiscent of CXCL12 binding pose in ACKR3^44^ (Fig. 2f). Moreover, the overall binding pocket in CCL7-DARC appears to be constricted compared to CCR2 (Extended Data Fig. 3c). A direct comparison of CCL7 positioning on DARC with all previously determined structures of chemokine-chemokine receptor complexes^45–50^ in active conformation further highlights the absence of CRS2 engagement in CCL7-DARC. As indicated in Fig. 2g, the N-terminus of CCL7 does not extend as deep into the orthosteric binding pocket as observed for other chemokines in complex with their cognate chemokine receptors. CCL7 displays maximal distance from the conserved tryptophan residue in TM6 (Trp^6.48^), bearing a close resemblance to CXCL12 interaction with ACKR3 (CXCR7) (Fig. 2g). In addition to CRS1, the 30s loop of CCL7 also forms a critical interaction interface by engaging the extracellular end of TM6 of the receptor (Fig. 2b).

**Fig. 2.**
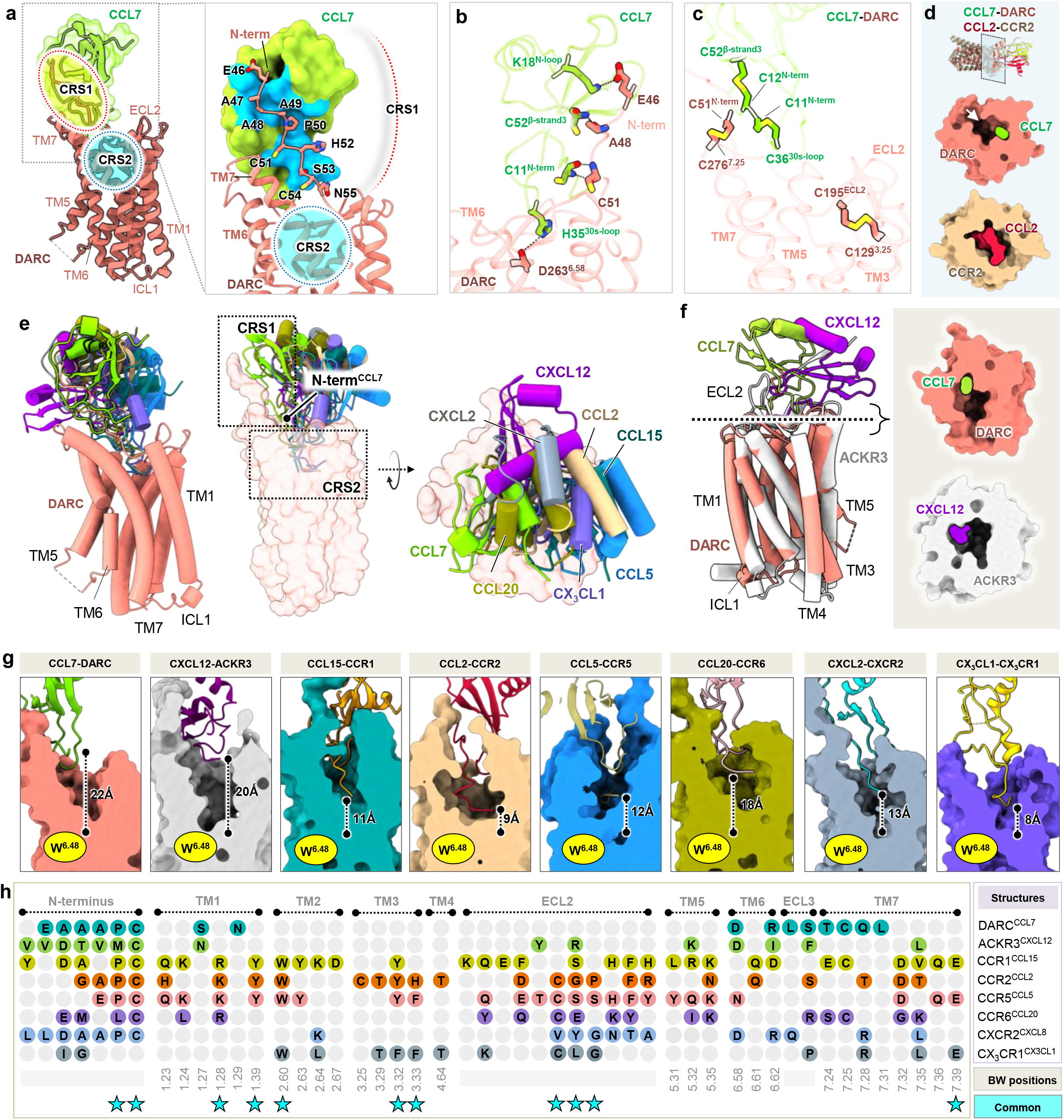
A distinct mode of chemokine recognition by DARC. **a.** Overall CCL7 binding mode in DARC. The regions of CRS1 and CRS2 are indicated by dashed circles (left). The key residues of DARC that interacts with CCL7 (blue surface) are shown. The possible CRS2 is empty and indicated by blue dashed circle. **b.** Interactions between the core domain of CCL7 and the N-terminus of DARC in CRS1 are illustrated (hydrogen bonds and salt-bridges are shown as black dashed lines). **c.** Disulfide linkages in CCL7 and DARC stabilize their overall architecture and fold. **d.** The shallow binding of CCL7 with DARC has been depicted as surface slice representation. The CCL2-CCR2 structure (PDB: 7XA3) has been used as a reference where CCL2 enters deep into the ligand binding pocket. **e.** Overall binding pose of chemokines on DARC (left and right) with CRS1 and CRS2 highlighted as dotted box (middle). Other receptor components have been removed after aligning the chemokine-receptor complex structures with CCL7-DARC. **f.** Structural alignment of CCL7-DARC and CXCL12-ACKR3 (left), the shallow binding mode of ligands on the respective ligands are illustrated as surface slice representation. **g.** Pairs of chemokine-chemokine receptor structures are shown to highlight the position of N-termini of chemokines at the orthosteric pocket. Distance between the N-terminus and the conserved toggle switch residue (Trp^6.48^) has been calculated in all structures. The N-terminus of CCL7 is the farthest from Trp^6.48^ compared to the other pairs (CXCL12-ACKR3, PDB: 7SK8; CCL15-CCR1, PDB: 7VL9; CCL2-CCR2, PDB: 7XA3; CCL5-CCR5, PDB: 7O7F; CCL20-CCR6, PDB: 6WWZ; CXCL8-CXCR2, PDB: 6LFO; CX_3_CL1-CX_3_CR1, PDB: 7XBX). **h.** Alignment of the interface residues between chemokines and their respective chemokine receptor structures have been provided. The Ballesteros-Weinstein positions are also mentioned below the TM regions. CCL7 makes the least number of contacts with DARC as compared to the other chemokine-chemokine receptor structures. The conserved interface residues in receptors are denoted as cyan colored stars.

We also compared the ligand-receptor contacts observed in CCL7-DARC with those in other chemokine-chemokine receptor complexes, and this further recapitulates a shallower binding mode of CCL7 to DARC. Importantly, the shallower binding of CCL7 with DARC translates to an absence of interactions between several residues in chemokines and TM1, TM2, TM3, TM4, ECL2, and TM5 observed in prototypical chemokine receptors (Fig. 2h). Thus, it is tempting to speculate that a superficial binding mode of chemokines to DARC, as observed here for CCL7, may impart chemokine promiscuity although it remains to be experimentally explored.

## Structural features of CCL7-bound DARC

In order to gain insights into overall structural features of CCL7-bound DARC, we compared our structure with the previously reported crystal structure of CCR2 in its inactive conformation^51, 52^ and cryo-EM structure of the same in its active conformation^46^. When compared to the inactive structure of CCR2 (PDB: 6GPS), TM5 and TM6 exhibit outward tilt angles of ∼35° and ∼12°, respectively (as measured from the Cα of Val219 for TM5 and Trp254 for TM6), while TM7 shows an inward helical tilt of ∼28° (as measured from the Cα of Thr300). However, while TM5 exhibits an outward tilt of ∼35° when compared to the CCL2 bound structure of CCR2 (PDB: 7XA3), TM6 and TM7 make inward tilt angles of ∼17° and ∼8°, respectively in the CCL7-bound structure of DARC (Fig. 3a). We also observed an overall similar pattern of TM movements when comparing the CCL7-DARC structure with inactive and active structures of CCR5^47, 53^ and CXCR2^54^ (Extended Data Fig. 4).

**Fig. 3.**
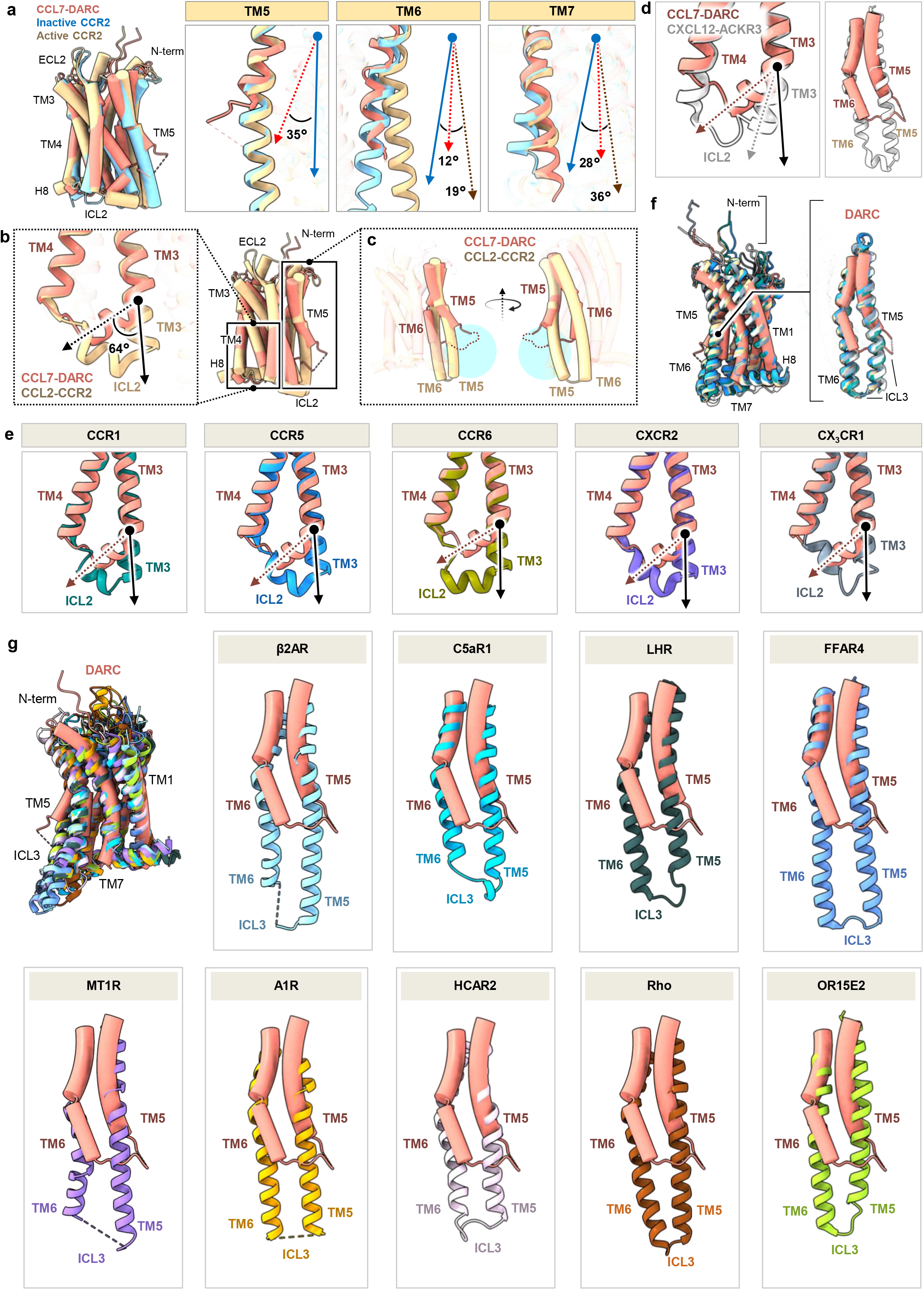
Divergent structural features of the 7TM domain in DARC. **a.** Structural alignment of CCL7-DARC with the inactive (PDB: 6GPS) and active (PDB: 7XA3) structures of CCR2 to highlight the changes in TM5, TM6 and TM7 (left). The deviation angles of TM5, TM6 and TM7 in CCL7-DARC and active CCR2 structure are shown against the inactive structure of CCR2. (Arrows with blue, red and brown depict the TMs of inactive CCR2, DARC and active CCR2, respectively). **b.** Conformations of ICL2 and TM4 were found to be unique in DARC as compared to the CCL2-CCR2 structure (PDB: 7XA3). The cytoplasmic end of TM3 forms a kink and translates about 64° in comparison to that of CCR2, probably due to a shorter ICL2. **c.** TM5 and TM6 are shorter in length compared to that of CCR2. In addition, TM5 exhibits an outward shift. **d.** TM3 in CXCL12-ACKR3 (PDB: 7SK8) forms a relatively smaller kink towards the cytoplasmic side when compared to CCL7-DARC (left). DARC harbors relatively shorter TM5 and TM6 as compared to ACKR3 (right). **e.** Conformations of ICL2 and TM4 in active chemokine upon alignment with CCL7 bound DARC. (CCL15-CCR1, PDB: 7VL9; CCL5-CCR5, PDB: 7O7F; CCL20-CCR6, PDB: 6WWZ; CXCL8-CXCR2, PDB: 6LFO; CX_3_CL1-CX_3_CR1, PDB: 7XBX). **f.** Superimposition of CCL7 bound DARC with active structures of chemokine-chemokine receptor pairs. Salmon, CCL7-DARC; gray, CXCL12-ACKR3 (PDB: 7SK8); blue, CCL5-CCR5 (PDB: 7O7F); beige, CCL2-CCR2 (PDB: 7XA3); deep gray, CXCL8-CXCR2 (PDB: 6LFO); teal, CCL15-CCR1 (PDB: 7VL9). DARC has been shown in cylinders and other structures are in ribbon representation. **g.** Relative lengths of TM5 and TM6 have been compared with active state classical GPCRs. A single receptor from each sub-family have been selected for comparison. (β2AR, PDB: 3SN6; C5aR1, PDB: 8HQC; LHR, PDB: 7FIG; FFAR4, PDB: 8G59; MT1R, PDB: 7VGZ; A1R, PDB: 6D9H; HCAR2, PDB: 7XK2; Rho, PDB: 6FUF; OR51E2, PDB: 8F76).

Interestingly, we observed that when compared to CCR2 the cytoplasmic portion of TM3 in DARC forms a kink at an angle of >60° at position Ala151^3.47^, while the intracellular loop 2 (ICL2) is relatively shorter (Fig. 3b). Strikingly, we also observed a significant shortening of TM5 and 6 in DARC compared to CCR2 although the ICL3 region spanning residues Gly227 to Asp238 is not resolved due to inherent flexibility (Fig. 3c). It is worth noting that the kink observed in TM3 of DARC is more pronounced than that observed in CXCR7, while the shortening of TM5 and 6 is not apparent in CXCR7 (Fig. 3d). Moreover, the TM3 kink is not observed in any of the previously determined active state structures of chemokine receptors, and therefore, it appears to be a unique feature of DARC (Fig. 3e). The striking shortening of TM5 and 6 also appears to be peculiar to DARC and is not observed in other chemokine receptors and prototypical GPCRs (Fig. 3f-g).

Next, we compared the conformational changes in the sequences corresponding to the conserved motifs present in prototypical GPCRs such as the DRY, CWxP, NPxxY and PIF motif. DARC lacks most of these motifs in its primary sequence, and instead harbors altered sequences in the corresponding regions, which display significantly different orientations compared to that observed in CCL2-bound CCR2 (Fig. 4a). It is interesting to note that similar to most chemokine receptors, DARC also contains a conserved P-C motif, constituted by Cys51 along with Pro50 in the N-terminus of the receptor. While the P-C motif in other chemokine receptors is located at the N-terminal loop immediately preceding TM1 and helps to bend the conformation of the N-terminal loop, the P-C motif in DARC adopts a linear conformation and is positioned away from TM1. The region of the N-terminus of DARC containing the P-C motif docks into a hydrophobic cavity in CCL7 wherein Pro50^N-term^ engages with Ile^42^ and Cys^52^ of CCL7, while Cys51^N-term^ interacts with Cys^11^ of CCL7 (Extended Data Fig. 5).

**Fig. 4.**
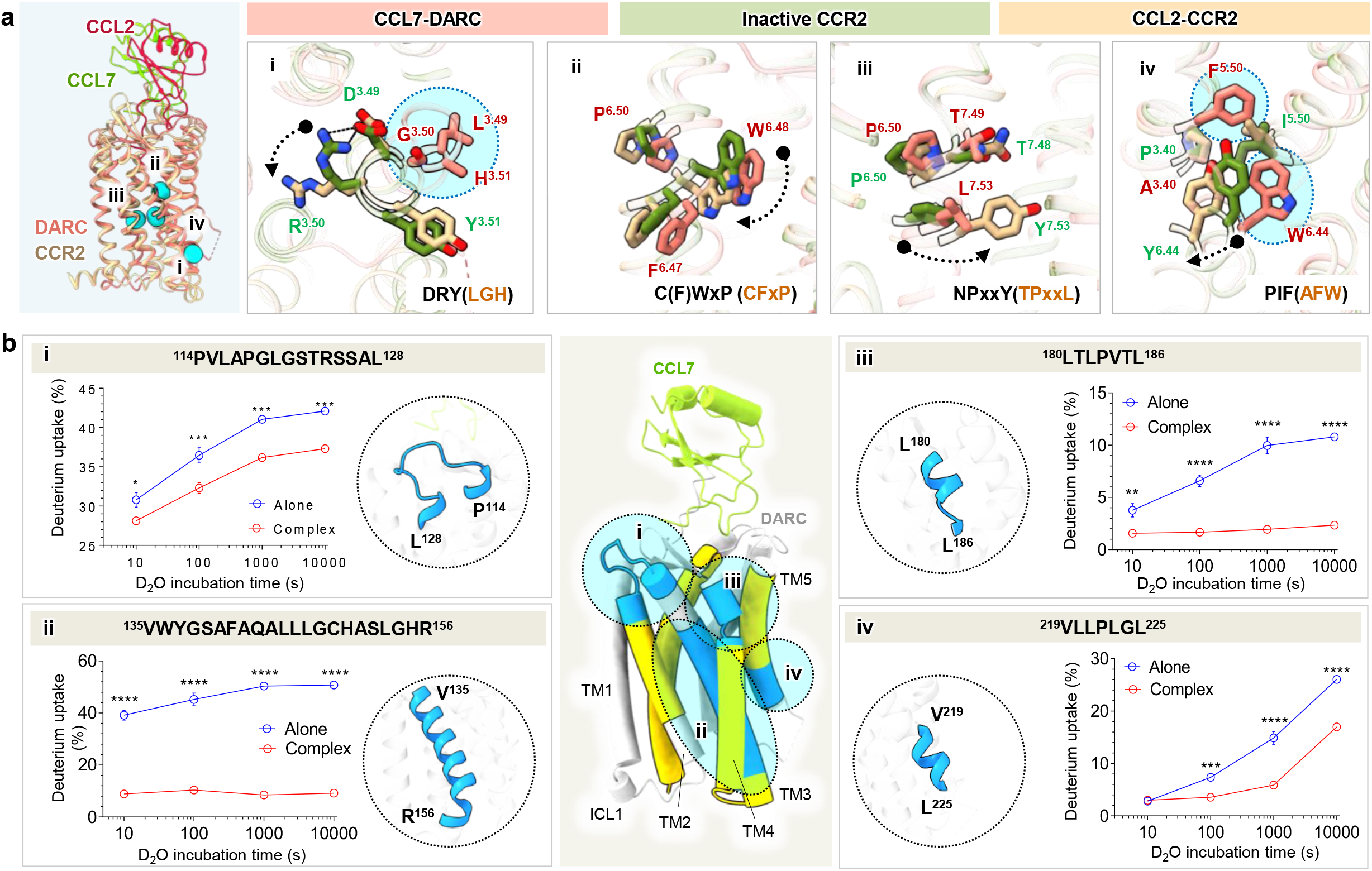
CCL7-induced conformational changes in DARC. **a.** The activation hallmark microswitches conserved in Class A receptors are shown in cyan spheres (left). Positions of microswitches are different in CCL7 bound DARC structure when compared to the classical GPCRs. The inactive (PDB: 6GPS) and active (PDB: 7XA3) structures of CCR2 have been taken as reference. **b.** HDX-MS profiles have been shown upon binding of CCL7 onto DARC. Different regions with distinct deuterium uptake values have been highlighted on the structure of DARC (blue, reduced deuterium exchange; yellow, no exchange). (a, b, c, d) The HDX-MS plots for the peptides with reduced exchange have been shown. Data (mean±SEM) represents three independent experiments analyzed using two-way ANOVA (Sidak’s multiple comparison; *p < 0.05, **p < 0.01, ***p < 0.001, ****p < 0.0001).

In order to measure CCL7-induced structural changes in DARC, we employed HDX-MS and compared the deuterium exchange for ligand-free and CCL7-bound receptor. While we observed limited coverage of the receptor as is typical with membrane proteins, we observed significant decrease in deuterium uptake in several regions of the receptor including ECL1, TM3, ECL2, and cytoplasmic end of TM5 upon binding of CCL7 (Fig. 4b and Extended Data Table 3). Considering that these regions are not directly involved in ligand binding, it likely reflects allosteric conformational changes in the receptor induced by binding of CCL7. Interestingly, the cytoplasmic ends of TM5 and 6, which exhibit maximal structural changes in prototypical GPCRs upon activation, do not appear to undergo major conformational changes in DARC, suggesting a distinct activation mechanism that remains to be experimentally determined.

## Structural basis of lack of canonical transducer-coupling

For prototypical GPCRs, including the chemokine receptors, agonist-induced activation results in a significant outward movement of TM5 and TM6 resulting in the formation of a cleft on the cytoplasmic side of the receptors (Fig.5a). This cytoplasmic cleft serves as the docking interface for the positioning of the α5 helix of G proteins^55, 56^, αN helix of GRKs^57^, and the finger loop of β-arrestins^58^, leading to stable coupling and subsequent activation. Specifically, this binding cleft on the receptor helps in mediating extensive interaction with the transducers through residues spanning TM3, TM5, TM6, ICL2 and ICL3^56^. The shortening of TM5 and 6, together with the distinct kink in TM3 and short ICL2, precludes the formation of such a binding cavity on the cytoplasmic side of DARC, thereby, presumably preventing the interaction with G proteins, GRKs and β-arrestins. As mentioned earlier, previous studies on DARC have been limited to the measurement of calcium response as a readout of G protein-coupling to DARC. Therefore, we first measured second messenger responses downstream of DARC, followed by a comprehensive profiling of all subtypes of G proteins using a previously described NanoBiT-based heterotrimer dissociation assay^59^. We observe that DARC fails to elicit any second messenger response upon stimulation by CCL7 (Fig. 5b-d), and dissociation of any of the G protein subtypes (Extended Data Fig. 6a). Similarly, DARC also does not exhibit any measurable coupling to any of the GRKs (Fig. 5e-g and Extended Data Fig. 6b-c) and β-arrestins (Fig. 5h-j and Extended Data Fig. 7). Considering the ligand promiscuity of DARC, we also measured cAMP response and β-arrestin recruitment following stimulation of the receptor by several C-C and C-X-C chemokines. However, similar to CCL7, we did not observe any detectable response for any of these chemokines in the transducer-coupling assays (Extended Data Fig. 8). Receptor surface expression was measured for all assays (Extended Data Fig. 9). Taken together, these data demonstrate how the 7TM fold in DARC has evolved to encode chemokine scavenging function in order to mediate chemokine homeostasis without canonical effector coupling and activation.

**Fig. 5.**
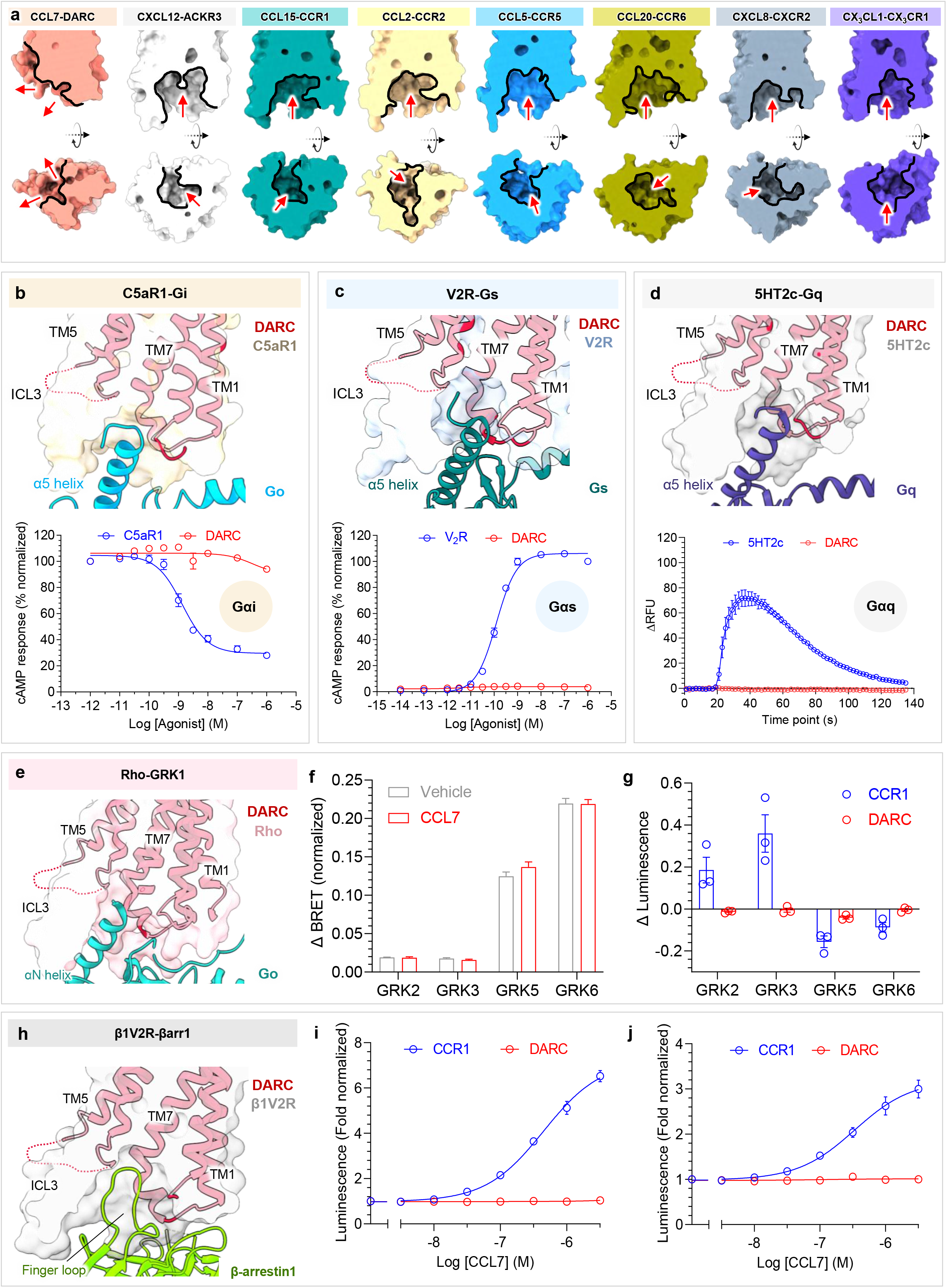
Molecular insights into the lack of canonical transducer-coupling to DARC. **a.** Superimposition of CXCL12-ACKR3 (PDB: 7SK8), CCL15-CCR1 (PDB: 7VL9), CCL2-CCR2 (PDB: 7XA3), CCL5-CCR5 (PDB: 7O7F), CCL20-CCR6 (PDB: 6WWZ), CXCL2-CXCR2 (PDB: 6LFO) and CX_3_CL1-CX_3_CR1 (PDB: 7XBX) with CCL7-DARC structure shown as surface slice representation illustrating the formation of the transducer binding cavity as opposed to CCL7-DARC. Red arrows highlight the formed cytoplasmic cavity. **b-d.** (top panel) C5aR1-Gi (PDB: 8HQC), V2R-Gs (PDB: 7BB6) and 5HT2c-Gq (PDB: 8DPF). DARC and G proteins are shown as cartoon and other receptors as transparent surface. (bottom panel) The G protein signaling profiles of DARC with respect to Go, Gi, Gs and Gq are shown in comparison with CCR1, C5aR1, V2R and 5HT2c, respectively. Data (mean±SEM) represents four (b-d) independent sets. Data has been normalized either with respect to the highest signal observed for each set (treated as 100%) (b) or with respect to the highest signal observed for positive control (treated as 100%) (c). **e.** Structural alignment of CCL7-DARC structure with Rho-GRK1 (PDB: 7MTB). **f, g.** Stimulation of DARC with CCL7 fails to induce GRK recruitment as measured by BRET assay (f) and NanoBiT assay (g). Data (mean±SEM) represents 3-6 independent experiments. Change in response has been plotted for both assays. **h.** Structural alignment of CCL7-DARC structure with β1V2R-βarr1 (PDB: 6TKO). **i, j.** Recruitment of βarr isoforms (βarr1/2) to DARC and CCR1 are shown, as assessed via NanoBit assay. Data (mean±SEM) represents three independent experiments and has been normalized with respect to the signal observed at basal condition for each set (treated as 1).

**Fig. 6.**
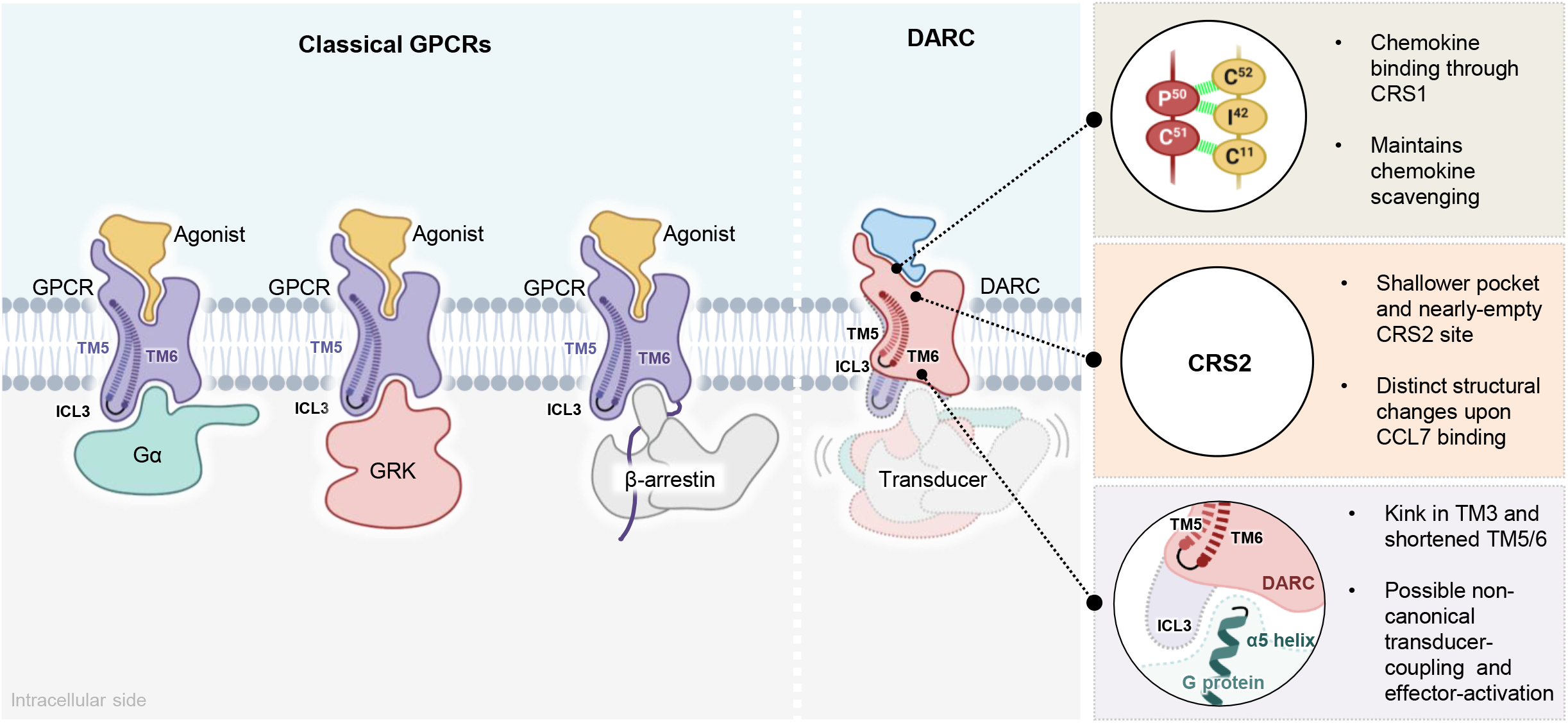
A schematic summary of distinct structural and functional features of DARC. Schematic representation highlighting ligand promiscuity and lack of transducer coupling of DARC in comparison to classical GPCRs. The schematic illustrates the importance of the cleft formed by the cytoplasmic ends of TM5 and TM6 in G protein, GRK and β-arrestin coupling (left). DARC exhibits the most promiscuous interaction with chemokines leading to scavenging functions. CRS2 in DARC is not directly involved in ligand binding. DARC possesses relatively shorter TM5/TM6 exhibiting limited conformational changes as compared to the canonical GPCRs, thereby precluding the interactions with G protein, GRK and β-arrestins (right).

## Concluding remarks

As mentioned earlier, CCL7 binds to DARC primarily through CRS1 without a significant involvement of CRS2. However, previous studies have proposed that in the two-site binding mechanism, CRS2 engagement by chemokines is important for allosterically directing transducer-coupling and effector-activation^43, 60^. Therefore, it is tempting to speculate that the absence of CRS2 engagement by chemokines upon binding to DARC, taken together with dramatic shortening of TM5/6, are the primary driving mechanisms underlying the functional divergence displayed by this receptor. However, the same remains to be established experimentally in future studies. While ACKRs are proposed to primarily serve chemokine scavenging functions^35, 36^, emerging evidence suggest their signaling capabilities in multiple cell types^40, 61–63^. Therefore, it would be important to probe the same for DARC in future studies, especially considering the recent data suggesting its expression in non-erythrocyte cell types and emerging indications of its fundamental role in several immune-physiological processes such as hematopoiesis^64^. While we used a full-length expression construct of DARC, the proximal N-terminus of the receptor is not resolved in the structure, and therefore, the quest to understand the interaction of PvDBP and PFTs to DARC remains open. In summary, we present a cryo-EM structure of CCL7-bound DARC, which elucidates the plausible molecular mechanism underlying chemokine scavenging and homeostasis even in the absence of canonical activation and signaling. This study should pave the way for further molecular and structural characterization of the intriguing functional divergence encoded in the 7TM scaffold exemplified by DARC and facilitate structure-guided design of novel therapeutics.

## ACKNOWLEDGMENTS

The research on the Duffy antigen receptor in A.K.S.’s laboratory was supported previously by the Intermediate Fellowship of the DBT Wellcome Trust India Alliance (IA/I/14/1/501285) and the Swarnajayanti Fellowship of the Department of Science and Technology (DST/SJF/LSA-03/2017-18), and currently by an MHRD-STARS grant (STARS/APR2019/BS/171/FS). The ongoing work in A.K.S.’s laboratory is also supported by the Senior Fellowship of the DBT Wellcome Trust India Alliance (IA/S/20/1/504916), Science and Engineering Research Board (SPR/2020/000408 and IPA/2020/000405), Indian Council of Medical research (F.NO.52/15/2020/BIO/BMS), Young Scientist Award from Lady Tata Memorial Trust, and IIT Kanpur. A.K.S. is an EMBO Young Investigator and Sonu Agrawal Memorial Chair. S.S. is funded by Prime Minister’s Research Fellowship (PMRF). We thank M. Baruah and S. Mahapatra for help with Nb52 purification, and M. Yadav for help with receptor purification. The work in V.M.K.’s laboratory was supported by a Swiss National Science Foundation Sinergia grant (grant 198545). HDX-MS work in K.Y.C.’s laboratory was supported by grants from the National Research Foundation of Korea funded by the Korean government (NRF-2021R1A2C3003518 and NRF-2019R1A5A2027340). A.I. was funded KAKENHI JP21H04791 and JP21H05113 from Japan by Society for the Promotion of Science (JSPS); JPMJFR215T and JPMJMS2023 from Japan Science and Technology Agency (JST); JP22ama121038 and JP22zf0127007 from the Japan Agency for Medical Research and Development (AMED). We also thank Kayo Sato, Shigeko Nakano and Ayumi Inoue in A.I.’s laboratory for their assistance in plasmid preparation and NanoBiT assays related to G-protein dissociation and GRK recruitment. The work in B.P.’s laboratory was supported a Wellcome Trust Seed Award in Science (215229/Z/19/Z), and Carole Daly was funded by a PhD studentship from the Department for the Economy (DfE), Queen’s University Belfast.

## AUTHOR CONTRIBUTIONS

SS expressed and purified ligand-receptor complexes, and carried out the GloSensor and βarr assays; BK prepared cryo-EM grids, optimized data collection, collected and processed cryo-EM under the supervision of VK; JM and RB processed cryo-EM data and determined the structure, JM prepared the figures with input from RB and SS; CM performed G-protein profiling and GRK interaction experiments using NanoBiT assay under the supervision of AI; CD and SH carried out BRET-based assay for GRK recruitment under the supervision of BP; HK performed the HDX-MS experiment under the supervision of KYC; PK carried out the calcium assay under the supervision of PNY; VK and AKS supervised the overall project and wrote the manuscript with input from all the authors.

## General reagents, plasmids, and cell culture

Most of the general reagents were purchased from Sigma Aldrich unless otherwise mentioned. Dulbecco′s Modified Eagle′s Medium (DMEM), Trypsin-EDTA, Fetal-Bovine Serum (FBS), Phosphate buffer saline (PBS), Hank’s balanced salt solution (HBSS), and Penicillin-Streptomycin solution were purchased from Thermo Fisher Scientific. HEK293T cells (ATCC) were maintained in DMEM (Gibco, Cat. no: 12800-017) supplemented with 10% (v/v) FBS (Gibco, Cat. no: 10270-106) and 100U/mL penicillin and 100µg/mL streptomycin (Gibco, Cat. no: 15140122) at 37°C under 5% CO_2_. *Sf9* cells were obtained from Expression Systems and maintained in protein-free cell culture media purchased from Expression Systems (Cat. no: 96-001-01) at 27°C with 135 rpm shaking. The cDNA coding region of DARC was cloned in pcDNA3.1 vector with an N-terminal FLAG tag and in pVL1393 vector with an N-terminal FLAG tag followed by the N-terminal region of M4 receptor (residue no. 2-23). All other receptor pcDNA3.1 constructs were also cloned with an N-terminal FLAG tag. The cDNA coding region of Nb52 was synthesized from Genscript, based on previously described sequence^42^ with an additional C-terminal 6X-His tag and cloned in pET22b(+) vector. All DNA constructs were verified by sequencing from Macrogen. For HDX-MS, a truncated version of DARC was used (27-322), cloned in pVL1393 vector with an N-terminal FLAG tag, followed by T4 lysozyme (T4L) and 3C protease cleavage site. Recombinant human C5a was purified from *E. coli* as described previously^65^. For the NanoBiT-G protein-dissociation assay and the flow cytometry-based expression analysis, the full-length human CCR1 or DARC was inserted into the pCAGGS expression vector with an N-terminal fusion of the hemagglutinin-derived signal sequence (ssHA), FLAG epitope tag and a flexible linker (MKTIIALSYIFCLVFA<u>DYKDDDDK</u>GGSGGGGSGGSSSGGG; the FLAG epitope tag is underlined), and the corresponding constructs were named as ssHA-FLAG-CCR1 and ssHA-FLAG-DARC, respectively. For the NanoBiT-β-arrestin-recruitment assay and the GRK-recruitment assay, ssHA-FLAG-CCR1 and ssHA-FLAG-DARC were C-terminally fused with the flexible linker and the SmBiT fragment GGSGGGGSGGSSSGG<u>VTGYRLFEEIL</u>; the SmBiT is underlined). The resulting plasmid was named as ssHA-FLAG-CCR1-SmBiT and ssHA-FLAG-DARC-SmBiT, respectively. Constructs of the NanoBiT-G protein sensors, the NanoBiT-β-arrestin sensors and the NanoBiT-GRK sensors were described previously^59, 66^.

## Purification of DARC

Full-length recombinant DARC was purified from *Sf9* insect cells as previously described for other GPCRs^67^. Briefly, *Spodoptera frugiperda* (*Sf9*) cells infected with DARC expressing baculovirus were harvested 72h post-infection. Cells were first homogenized in hypotonic buffer (20mM HEPES pH 7.4, 20mM KCl, 10mM MgCl₂, 1mM PMSF, 2mM benzamidine) followed by hypertonic buffer (20mM HEPES pH 7.4, 20mM KCl, 10mM MgCl_2_, 1M NaCl, 1mM PMSF, 2mM benzamidine). Cells were then lysed by solubilization in lysis buffer (20mM HEPES pH 7.4, 450mM NaCl, 1mM PMSF, 2mM benzamidine, 0.1% cholesteryl hemisuccinate, 2mM iodoacetamide and 1% L-MNG) for 2h at 4°C. The lysate was then diluted in buffer containing 20mM HEPES pH 7.4, 2mM CaCl₂, 1mM PMSF, and 2mM benzamidine and cleared by centrifuging at 20,000 rpm for 30mins. The supernatant was collected, filtered and loaded onto pre-equilibrated M1-FLAG column. Following loading, the column was washed alternatively with low salt buffer (20mM HEPES pH 7.4, 150mM NaCl, 2mM CaCl₂, 0.01% cholesteryl hemisuccinate, 0.01% L-MNG) and high salt buffer (20mM HEPES pH 7.4, 350mM NaCl, 2mM CaCl₂, 0.01% L-MNG). Protein was eluted using 250μg/mL FLAG and 2mM EDTA. 100nM CCL7 was kept in all buffers throughout the purification process. Free cysteines on the receptor surface were blocked with iodoacetamide, and excess iodoacetamide was quenched using free L-cysteine. Purified receptor-ligand complex was stored in −80°C till further use.

For HDX-MS, DARC was purified in the absence of CCL7. Following receptor blocking with iodoacetamide, the sample was passed through PD-10 desalting columns to remove excess FLAG, iodoacetamide and L-cysteine from the solution. The receptor was then incubated with 3C-protease (to cleave the N-terminal FLAG and T4L tag) and PNGase (to remove glycosylation). The sample was then concentrated and injected into size-exclusion chromatography (SEC) to isolate pure cleaved receptor. For CCL7-DARC complex, cleaved DARC was incubated with 1.5 molar excess of CCL7 prior to SEC. Samples were stored with 10% glycerol in −80°C till further use.

## Purification of CCL7

Recombinant human CCL7 was purified from *E. coli* as described previously with slight modifications^68^. Briefly, *E. coli* cells transformed with cDNA region of CCL7 (containing an N-terminal 6X-His tag followed by enterokinase cleavage site) cloned in pGEMEX vector were cultured at 27°C till O.D._600_ of 1.5 and induced with 1mM IPTG. This was followed by an additional culturing of 48h at 20°C. Cell pellet thus obtained was resuspended in lysis buffer (20mM HEPES pH 7.4, 1M NaCl, 10mM imidazole, 5% glycerol, 0.3% Triton-X-100 and 1mM PMSF) and lysed by sonication for 20mins. The lysate was centrifuged at 18,000 rpm for 30mins to remove cellular debris and the supernatant was filtered and loaded onto pre-equilibrated Ni-NTA beads. After loading, the beads were washed with wash buffer (20mM HEPES pH 7.4, 1M NaCl and 40mM imidazole) and bound protein was eluted using elution buffer (20mM HEPES pH 7.4, 1M NaCl and 500mM imidazole). The eluted protein was dialyzed overnight at 4°C against dialysis buffer (50mM Tris pH 8.0 and 50mM NaCl) to remove imidazole and to bring the protein in a buffer compatible with enterokinase activity. This was followed by enterokinase treatment and subsequent cation-exchange chromatography to separate the cleaved protein from un-cleaved protein. Fractions corresponding to the protein of interest were pooled, dialyzed overnight at 4°C against 20mM HEPES pH 7.4 and 150mM NaCl, and stored with 10% glycerol in −80°C till further use.

## Purification of Nb52

Nb52 was purified from *E. coli Rosetta* strain. Briefly, cells were cultured in 2XYT media at 37°C and induced with 100µM IPTG at an O.D._600_ of 0.6, followed by culturing for an addition 18h at 18°C. Cells were resuspended in lysis buffer (20mM HEPES pH 8.0, 150mM NaCl, 1mM PMSF, 2mM benzamidine and 1mg/mL lysozyme) and lysed by sonication for 20mins. The lysate was centrifuged at 18,000 rpm to clear the cellular debris, and the resulting supernatant was filtered and loaded onto pre-equilibrated Ni-NTA beads. Following loading, the beads were washed with wash buffer (20mM HEPES pH 8.0, 150mM NaCl and 30mM imidazole) and bound protein was eluted with elution buffer (20mM HEPES pH 8.0, 150mM NaCl and 500mM imidazole). The eluted protein was subjected to size-exclusion chromatography. Fractions corresponding to the protein of interest were pooled and stored with 10% glycerol in −80°C till further use.

## Negative staining electron microscopy

Conventional uranyl formate negative staining was performed for judging sample homogeneity prior to grid freezing for cryo-EM following the protocol described earlier^69^. Briefly, 3.5µl of the sample was dispensed onto a freshly glow discharged formvar/carbon coated grid (*Ted Pella*), incubated for 1min and blotted off using a Whatman No. 1 filter paper. The grid with the adhered sample was touched onto a first drop of freshly prepared 0.75% uranyl formate stain, and immediately blotted off using a filter paper. The grid was then touched onto a second drop of uranyl formate and moved in a rotating fashion for 30s to increase the efficiency of staining. The excess stain was blotted-off and air dried prior to imaging and data collection. Imaging and data collection were performed with a FEI Tecnai G2 12 Twin TEM (LaB6) equipped with a Gatan 4k x 4k CCD camera at 30,000x magnification and operating at 120kV. The collected micrographs were imported into Relion 3.1.2^70–72^ for subsequent processing. Approximately 10,000 particles were automatically picked using gaussian blob picker, extracted with a box-size of 280px and subjected to reference free 2D classification to obtain the 2D class averages.

## Cryo-EM data collection and acquisition

3.5µl of the protein sample at a concentration of 3-4mg/mL was applied onto a Quantifoil 1.2/1.3 grid (300 mesh) glow discharged for 25s using a PELCO easiGlow (*Ted Pella*) cleansing system. The grid was blotted for 3s using Whatman no. 1 filter papers and flash frozen in liquid ethane (−181°C) with a Vitrobot MarkIV (Thermo Fisher Scientific) maintained at 4°C and 100% humidity. Data collection was performed on a 300kV Titan Krios (Thermo Fisher Scientific) equipped with a Gatan K3 direct electron detector in super-resolution mode. Movie stacks consisting of 40 frames were collected automatically using EPU at a nominal magnification of 130,000x and a pixel size of 0.65Å over a defocus range of 1-2µm with a total dose of 58e^-^/Å^2^ and 80e^-^/Å^2^ for the first and second datasets, respectively.

## Cryo-EM data processing

17,287 dose-fractionated movies were collected over two independent sessions. First, all these movies were assigned to distinct optics groups according to the EPU beam shift values using a script provided by Dr. Pavel Afanasyev (ETH Zurich; https://github.com/afanasyevp/afis). Subsequently, the movies were subjected to beam-induced motion correction in Relion 3.1.3 and Relion 4.0. All processing steps hereafter were performed with cryoSPARC v4.0^73^. The motion corrected micrographs were imported into cryoSPARC v4.0 and CTF parameters were estimated with Patch CTF (Multi) followed by automated particle picking with the blob-picker sub-program yielding 12,570,207 particles. These auto-picked particles were extracted with a box size of 416px (fourier cropped to 64px) and subjected to several rounds of reference free 2D classification. 2D class averages with clear secondary features representing all possible orientations were selected and used as templates for template-based particle picking. 10,833,233 particles from the template picker job were extracted with a box size of 416px (fourier cropped to 64px) and subjected to multiple rounds of 2D classification. The particles corresponding to the clean 2D class averages from both the blob-picker and template picker jobs were selected, combined, and duplicate particles were removed with the “Remove duplicates” sub-program. The resulting particle stack was re-extracted with a box size of 416px (fourier cropped to 288px) and subjected to ab-initio reconstruction, yielding 3 classes. The best class containing 313,511 particles was subjected to local refinement with a mask on the dimeric complex, yielding a reconstruction with 4.3Å global resolution. Imposing C2 symmetry during subsequent local refinements with mask on the dimeric complex led to an improvement in resolution to 3.9Å and helped to resolve the side chains. In order to boost the resolution further, particles corresponding to the micrographs with contrast transfer function (CTF) fit better than 3.5Å were selected and re-extracted with full box size of 416px. Subsequent local refinement with these 240,185 particles led to a map with global resolution of 3.8Å at 0.143 FSC cut-off. The detailed processing pipeline has been included as Extended Data Fig. 2. Local resolution estimation of the final refined map was performed with the Blocres sub-program, while map sharpening was done with the “Sharpening tools” sub-module within the cryoSPARC suite.

## Model building and validation

The starting coordinates for DARC was derived from an AlphaFold model (AF-Q4VBN9-F1-model_v4)^74^ while the coordinates of CCL7 was obtained from a previously solved crystal structure of chemokine binding protein of orf virus complexed with CCL7 (PDB: 4ZKC)^75^. These initial models were used to dock into the coulombic map of DARC-CCL7 complex with Chimera^76, 77^ to generate the starting dimeric coordinates. The dimeric model so obtained was subjected to flexible fitting with the “all-atom refine” sub-module in COOT^78^, followed by iterative rounds of manual readjustments of the side chains in COOT and refinement of the rebuilt model against the EM map with phenix.real_space_refine^79, 80^. The final refined model had 93.57% of the residues in most favored region and 6.53% in the allowed region of the Ramachandran plot and were validated with Molprobity^81^. Data collection, processing and model refinement statistics are provided as Extended Data Table 1. Figures included in the manuscript have been prepared with Chimera or ChimeraX software.

## Hydrogen-Deuterium Exchange Mass Spectrometry

CCL7, DARC and CCL7-DARC complex were prepared at 71µM. 4.5µl of the protein samples were mixed with 25.5µl of D_2_O buffer (20mM HEPES pD 7.4, 150mM NaCl) and incubated for 10, 100, 1000, and 10,000s at room temperature. The deuterated samples were quenched by adding 30µl of ice-cold quench buffer (0.1M NaH_2_PO4 pH 2.01, 20mM TCEP and 10% glycerol) and snap-frozen on dry ice and stored at −80°C. Non-deuterated samples were prepared by mixing 4.5µl of protein samples with 25.5µl of their respective H_2_O buffers, followed by the same quenching and freezing steps, as described above. The quenched samples were digested and isolated using the HDX-UPLC-ESI-MS system (Waters, Milford, MA, USA). The quenched samples were thawed and immediately passed through an immobilized pepsin column (2.1x30mm) (Life Technologies, Carlsbad, CA, USA) at a flow rate of 50µl/min in 0.05% formic acid in H_2_O at 12°C. The peptic peptides were collected on a C18 VanGuard trap column (1.7µmx30mm) (Waters) for desalting with 0.05% formic acid in H_2_O and then separated by an Acuity UPLC C18 column (1.7µm, 1.0x100mm) (Waters) at a flow rate of 40µl/min with an acetonitrile gradient starting from 8% B and increasing to 85% B over 8.5mins. The mobile phase A was 0.1% formic acid in H_2_O, and the mobile phase B was 0.1% formic acid in acetonitrile. Buffers were adjusted to pH 2.5 and the system was maintained at 0.5°C (except pepsin digestion which was at 12°C) to minimize back-exchange of deuterium. Mass spectral analyses were performed by Xevo G2 quadrupole-time of flight (Q-TOF) equipped with a standard ESI source in MSE mode (Waters) with positive ion mode. The capillary, cone, and extraction cone voltages were set to 3kV, 40V, and 4V, respectively. Source and desolvation temperatures were set to 120°C and 350°C, respectively. Trap and transfer collision energies were set to 6V, and trap gas flow rate was set to 0.3mL/min. Sodium iodide (2µg/µl) was utilized to calibrate the mass spectrometer and [Glu1]-Fibrinopeptide B (200fg/µl) in MeOH:water (50:50 (v/v) + 1% acetic acid) was utilized for lock-mass correction. The ions at mass-to-charge ratio (m/z) 785.8427 were monitored at scan time 0.1s with a mass window of ±0.5Da. The reference internal calibrant was introduced at a flow rate of 20µl/min and all spectra was automatically corrected using lock-mass. Two independent interleaved acquisition functions were created: the first function, typically set at 4eV, collected low energy or unfragmented data whereas the second function collected high energy or fragmented data typically obtained by using a collision ram form 30-55eV. Ar gas was used for collision induced dissociation (CID). Mass spectral were acquired in the range of m/z 100-2000 for 10mins. ProteinLynx Global Server 2.4 (Waters) was utilized to identify peptic peptides from the non-deuterated samples with variable methionine oxidation modification and a peptide score of 6. DynamX 3.0 (Waters) was used to determine the level of deuterium uptake for each peptide by measuring the centroid of isotopic distribution. Detailed HDX-MS data information is provided in Extended Data Table 3.

## Second messenger signaling assays

To measure the effect of ligand stimulation on Gs and Gi-mediated signaling, intracellular cAMP levels were measured using the GloSensor assay, as previously described^40^. In brief, HEK293T cells were transfected with either 3.5μg of FLAG-tagged DARC/V_2_R or 2.5μg of FLAG-tagged C5aR1 (encoded in pcDNA vector) and 3.5μg of F22 plasmid (Promega, Cat. no: E2301). 14-16h following transfection, cells were trypsinized and seeded in 96-well plates at a density of 100,000 cells/well in assay buffer (20mM HEPES pH 7.4, 1X HBSS and 0.5mg/mL D-luciferin (GoldBio, Cat. no: LUCNA-1G)) and incubated for 1h 30mins at 37°C followed by an additional 30mins at room temperature. Basal luminescence was measured for 5 cycles. For measuring Gi-mediated decrease in cytosolic cAMP levels, 5μM forskolin was added to the wells and luminescence was measured for 8 cycles till the reading stabilized. This was followed by addition of ligand at the indicated final concentration. Luminescence was recorded for 30 cycles. Signal obtained was normalized either with respect to reading observed for positive control at the highest concentration of ligand (for measuring Gs-mediated increase in intracellular cAMP levels) or with signal observed at the lowest concentration of ligand for each receptor (for measuring Gi-mediated decrease in intracellular cAMP levels), treated as 100%.

Calcium flux assay was undertaken to measure ligand induced changes in cytosolic Ca^2+^ levels, as previously described^41^. Briefly, HEK293T cells were transfected with 4μg of pGP-CMV-GcAMP6s (Ca^2+^ sensor plasmid; Addgene, Cat. no: 40753) and either 4μg of 5HT_2c_ receptor (as positive control) or 4μg of DARC using PEI max in a ratio of 1:4 (DNA:PEI max). Transfected cells were seeded in a black optical bottom plate at a density of 50,000 cells/well in complete medium (supplemented with 10% FBS). 14-16h post-transfection, media from the wells was replaced with 100μL of Ca^2+^/Mg^2+^ free HBSS buffer (pH 7.2) and the cells were incubated for an additional 10mins at 37°C in Flex Station 3 (Molecular Devices). Ligand induced changes in the relative fluorescence unit (RFU) was measured at an excitation wavelength of 485nm and emission wavelength of 525nm (cut off 515nm) with a setting of 6 reads/well. Basal fluorescence was recorded for 15s for each well, followed by addition of 20μL of 6X concentration of ligand using robotic pipetting of FlexStation system. RFU was recorded at an interval of 2s for a total duration of 135s. The change in RFU (ΔRFU) for each group was calculated by subtracting the average basal response (RFU before ligand addition) from RFU of each well at each time point following ligand addition. ΔRFU was plotted and analyzed using GraphPad Prism 9 software.

## NanoBiT-based G protein dissociation assay

Ligand-induced G protein activation was measured using a previously described NanoBiT-based G protein dissociation assay^59^. Briefly, a NanoBiT-G protein consisting of LgBiT-tagged Gα subunit and SmBiT-tagged Gγ2 subunit along with the untagged Gβ1 subunit were co-expressed with the indicated receptor constructs and ligand-induced change in luminescence signal was measured. Typically, HEK293 cells (Thermo Fisher Scientific) were seeded in a 6-well culture plate and transfected with a plasmid mixture consisting of 100ng LgBiT-Gα (Gαs, Gαi1, Gαi2, Gαi3, Gαo, Gαq, Gα12 or Gα13), 500ng Gβ1, 500ng SmBiT-Gγ2 (C68S) with 200ng receptor plasmid (ssHA-FLAG-CCR1, ssHA-FLAG-DARC or an empty plasmid). To enhance NanoBiT-G protein expression for Gs, Gq and G12/13, 100ng of RIC8B plasmid (isoform 2; for Gs) or RIC8A (isoform 2; for Gq, G12, and G13) were co-transfected. After 24h of transfection, the cells were harvested with EDTA-containing PBS, centrifuged, and suspended in 2mL of Hank’s Balanced Salt Solution (HBSS) containing 0.01% bovine serum albumin (BSA fatty acid–free grade, SERVA) and 5mM 4-(2-hydroxyethyl)-1-piperazineethanesulfonic acid (HEPES) at pH 7.4 (assay buffer). Afterwards, the cells were dispensed in a white 96-well plate (80μL/well), incubated with 20μL of 50μM coelenterazine (Carbosynth, Cat. no: EC175526), and 2h later, baseline luminescence was measured (SpectraMax L, Molecular Devices). Subsequently, 20μL of 6X CCL7, serially diluted in the assay buffer, was manually added and the plate was immediately read for the second measurement in kinetic mode. Luminescence counts recorded from 5 to 10min post-agonist addition were averaged, corrected with the baseline signal, normalized with respect to vehicle control and plotted using the GraphPad Prism 9 software.

## NanoBiT-based β-arrestin/GRK recruitment assay

Ligand-induced recruitment of β-arrestin or GRK was performed as described previously with minor modifications^66^. Transfection was performed according to the same procedure as described in the “NanoBiT-based G protein-dissociation coupling assay” section except for a plasmid mixture consisting of 500ng ssHA-FLAG-GPCR-SmBiT and 100ng LgBiT-β-arrestin (β-arrestin assay) or 500ng ssHA-FLAG-GPCR-SmBiT and 500ng GRK-LgBiT (GRK assay). The transfected cells were dispensed into a 96-well plate, and ligand-induced luminescent changes were measured following the same procedures as described for the NanoBiT-based G protein-dissociation coupling assay.

## Bioluminescence Resonance Energy Transfer (BRET)-based GRK recruitment assay

HEK293 clonal cell line (HEK293SL cells), referred to as HEK293 cells, were a gift from Stephane Laporte (McGill University, Montreal, Quebec, Canada). These cells were cultured in Dulbecco’s Modified Eagle’s Medium (DMEM) high glucose (Gibco) supplemented with 10% fetal bovine serum and 100units/mL penicillin-streptomycin (Gibco), maintained at 37°C and 5% CO_2_ and passaged every 3-4 days using trypsin-EDTA 0.05% (Gibco) to detach the cells. DNA to be transfected was combined with salmon sperm DNA (Invitrogen) to obtain a total of 1.6µg DNA per condition. Increasing amounts of GFP10-GRK (0 to 800ng) was transfected along with 100ng of Gβ1, 100ng of Gγ2 and 50ng of DARC-RlucII. Linear polyethyleneimine 25K (PEI; Polysciences) was combined with DNA (4.8µg PEI per 1.6µg of DNA), vortexed and incubated for 20mins before adding 1.2mL of cell suspension containing 360,000 cells for each condition. Cells containing the DNA were seeded (100μL/well) in white 96-well plates (Greiner) and incubated for 48h before assay. The day of the assay, cells were washed with DPBS (Gibco) and assayed in Tyrode’s buffer (137mM NaCl, 0.9mM KCl, 1mM MgCl_2_, 11.9mM NaHCO_3_, 3.6mM NaH_2_PO_4_, 25mM HEPES, 5.5mM glucose, 1mM CaCl_2_ (pH 7.4)) at 37°C. The expression of GFP10 was monitored in each well using a FLUOstar Omega microplate reader (BMG Labtech) by exciting at 400nm and reading the fluorescence at 520nm. CCL7 100nM final or vehicle were added and cells were incubated at 37°C for 30mins. 5mins before reading, 2.5µM of the Renilla luciferase substrate (coelenterazine 400a; NanoLight Technology) was added. BRET measurements were performed using the FLUOstar Omega microplate reader with an acceptor filter (515±30nm) and donor filter (410±80nm). BRET was calculated by dividing GFP10 emission by RlucII emission.

## Co-immunoprecipitation to detect physical interaction between DARC and β-arrestin1/2

Co-immunoprecipitation (coIP) was performed to measure the physical coupling between DARC and β-arrestin1/2, as described earlier^82^. In brief, HEK293T cells transfected with 3.5μg of FLAG-tagged receptor and either 1.7μg of β-arrestin1 or 3.5μg of β-arrestin2 were harvested 48h post-transfection. Prior to harvesting, cells were serum starved for 6h and stimulated with 100nM of CCL7. The pellets were dounced in buffer containing 20mM HEPES pH 7.4, 450mM NaCl, 1X PhosSTOP (Sigma, Cat. no: 4906845001) and 1X cOmplete protease inhibitor cocktail (Sigma, Cat. no: 4693116001), and incubated with 1.5mM freshly prepared DSP (Sigma, Cat. no: D3669) for 45mins at room temperature under tumbling conditions. This was followed by solubilization, wherein the suspension was incubated with 0.1% L-MNG for 1h at room temperature under tumbling conditions. Cell debris was removed by centrifugation, and the supernatant was incubated with pre-equilibrated anti-FLAG M1 beads for 1h 30mins, in the presence of 2mM CaCl_2_, at room temperature. Beads were washed five times alternatively with low salt buffer (20mM HEPES pH 7.4, 150mM NaCl, 2mM CaCl₂, 0.01% CHS, 0.01% L-MNG) and high salt buffer (20mM HEPES pH 7.4, 350mM NaCl, 2mM CaCl₂, 0.01% L-MNG). Elution was collected by incubating the beads with elution buffer (20mM HEPES pH 7.4, 150mM NaCl, 2mM EDTA, 0.01% L-MNG, 250μg/μL FLAG) for 30mins. Eluted samples were run on a SDS-PAGE followed by western blotting. The blots were blocked with 1% BSA for 1h at room temperature, followed by incubation with rabbit anti-βarr antibody (1:2500; CST, Cat. no: 4674) overnight at 4°C. The next day, the blots were washed thrice with 1X TBST (10mins each wash) and then incubated with anti-rabbit secondary antibody (1:10,000) for 1h at room temperature. The blots were once again washed three times with 1X TBST (10mins each wash) and developed with Promega ECL solution on a chemidoc (BioRad). The blots were then stripped and re-probed for receptor using anti-FLAG M_2_HRP (1:5000, Sigma, Cat. no: A8592). Bands were quantified using ImageLab software and normalized with respect to signal obtained at highest time point for positive control (treated as 100%). To assess whether addition of V_2_-tail can promote recruitment of β-arrestin, several chimeric constructs were generated wherein the V_2_-tail was added after truncating the C-terminal end of DARC to different extents. For all of these chimeric N-terminal FLAG-tagged receptor constructs, 3.5μg of DNA was transfected.

## Tango Assay to measure β-arrestin2 recruitment

To measure β-arrestin2 recruitment following ligand stimulation, Tango assay was undertaken^83, 84^. Briefly, HTLA cells were transfected with 7μg of FLAG-tagged receptor. 24h post-transfection, cells were trypsinized and seeded in 96-well plates at a density of 100,000 cells/well in complete media and allowed to adhere. Next day, complete media was replaced with incomplete DMEM supplemented with respective ligand at indicated dose. After 8h, incomplete media was removed from the wells and drug buffer (20mM HEPES pH 7.4, 1X HBSS, 0.5mg/mL D-luciferin) was added. Emitted luminescence was recorded immediately. Signal was normalized with respect to reading observed for positive control at the highest concentration dose, treated as 100%.

## Confocal microscopy to visualize β-arrestin1/2 trafficking

To visualize β-arrestin recruitment to the cell membrane and subsequent endocytosis, confocal assay was performed as described earlier^85^. HEK293 cells transfected with 3.5μg of FLAG-tagged receptor and 3.5μg of YFP-tagged β-arrestin1/2 were seeded in confocal dishes (pre-coated with 0.01% poly-D-Lysine) at a density of 1 million cells/dish, 24h post-transfection. Cells were allowed to adhere to the confocal dishes, and 24h post-seeding, cells were serum-starved for 6h. Confocal imaging of all samples was done using Zeiss LSM 710 NLO confocal microscope wherein samples were housed on a motorized XY stage with a CO_2_ enclosure and a temperature-controlled platform equipped with 32x array GaAsP descanned detector (Zeiss). A multi-line argon laser source was used for the green channel (mYFP). All microscopic settings including pinhole opening and laser intensity were kept in the same range for a parallel set of experiments. Cells were stimulated with 100nM CCL7 for indicated time duration. Images were scanned in line scan mode and acquired images were processed post imaging in ZEN lite (ZEN-blue/ZEN-black) software suite from ZEISS.

## Receptor surface expression assay

To assess cell surface expression of the corresponding receptors, whole cell surface ELISA was performed, as previously described^86^. Briefly, transfected cells were seeded at a density of 0.1 million cells/well 24h post-transfection in 24-well plates (pre-coated with 0.01% poly-D-Lysine) and incubated for 24h at 37°C in a CO_2_ incubator. Once cells were well adhered, the media was removed by aspiration and the cells were washed once with 1X TBS. This was followed by treatment with 300μL 4% PFA for 20mins to fix the cells. Excess PFA was removed by washing the cells thrice with 400μL of 1X TBS. The wells were blocked by incubating with 200μL 1% BSA for 1h 30mins. 200μL of anti-FLAG M_2_HRP antibody (Sigma, Cat no. A8592) at a dilution of 1:10,000 was added to the wells and incubated for an additional 1h 30mins. To develop the signal, 200μL of TMB (Thermo Scientific, Cat. no: 34028) was added to the wells and incubated till the development of adequate color. Reaction was quenched by transferring 100μL of solution to a 96-well plate containing 100μL of 1M H_2_SO_4_ and absorbance was measured at 450nm. To estimate the amount of cells fixed in each well, excess TMB was first removed by washing once with 400μL of 1X TBS and the cells were incubated for 20mins in 0.2% Janus green B (Sigma, Cat. no. 201677). After removing excess stain by repeated washing with MQ water, signal was developed by adding 800μL of 0.1N HCl to each well and read at 595nm. Signal was normalized by dividing the reading obtained at 450nm with the reading obtained at 595nm. Receptor surface expression was fold normalized with respect to vehicle (empty pcDNA vector) transfected cells.

For the NanoBiT-based G protein dissociation assay, surface expression of the receptors was measured using flow-cytometry based assay following a previously described protocol^40^. HEK293 cells were transfected with plasmids for ssHA-FLAG-CCR1 or ssHA-FLAG-DARC along with the NanoBiT-Gi1 sensor, as described in the “NanoBiT-based G protein-dissociation assay” section. The cells were harvested with 0.5mM EDTA-containing PBS and transferred to a 96-well V-bottom plate. The cells were then fluorescently labelled using anti-FLAG monoclonal antibody (Clone 1E6, FujiFilm Wako Pure Chemicals; 10μg/mL diluted in 2% goat serum and 2mM EDTA-containing PBS) followed by incubation with Alexa Fluor 488-conjugated goat anti-mouse IgG secondary antibody (Thermo Fisher Scientific; 10μg/mL). Subsequently, the cells were washed with PBS, resuspended in 2mM EDTA-containing PBS, filtered through a 40μm filter and the fluorescent intensity of single cells was quantified using a flow cytometer. Fluorescent signal from Alexa Fluor 488 was recorded and analyzed using the FlowJo software. Mean fluorescence intensity from about 20,000 cells per sample were used for analysis. Typically, we obtained a CCR1 MFI value of 2,000 (arbitrary unit) and a mock MFI value of 20. For each experiment, we normalized an MFI value of DARC by that of CCR1 performed in parallel and denoted relative expression levels.

