## Supplemental Material for "Structure of the human Duffy antigen receptor"

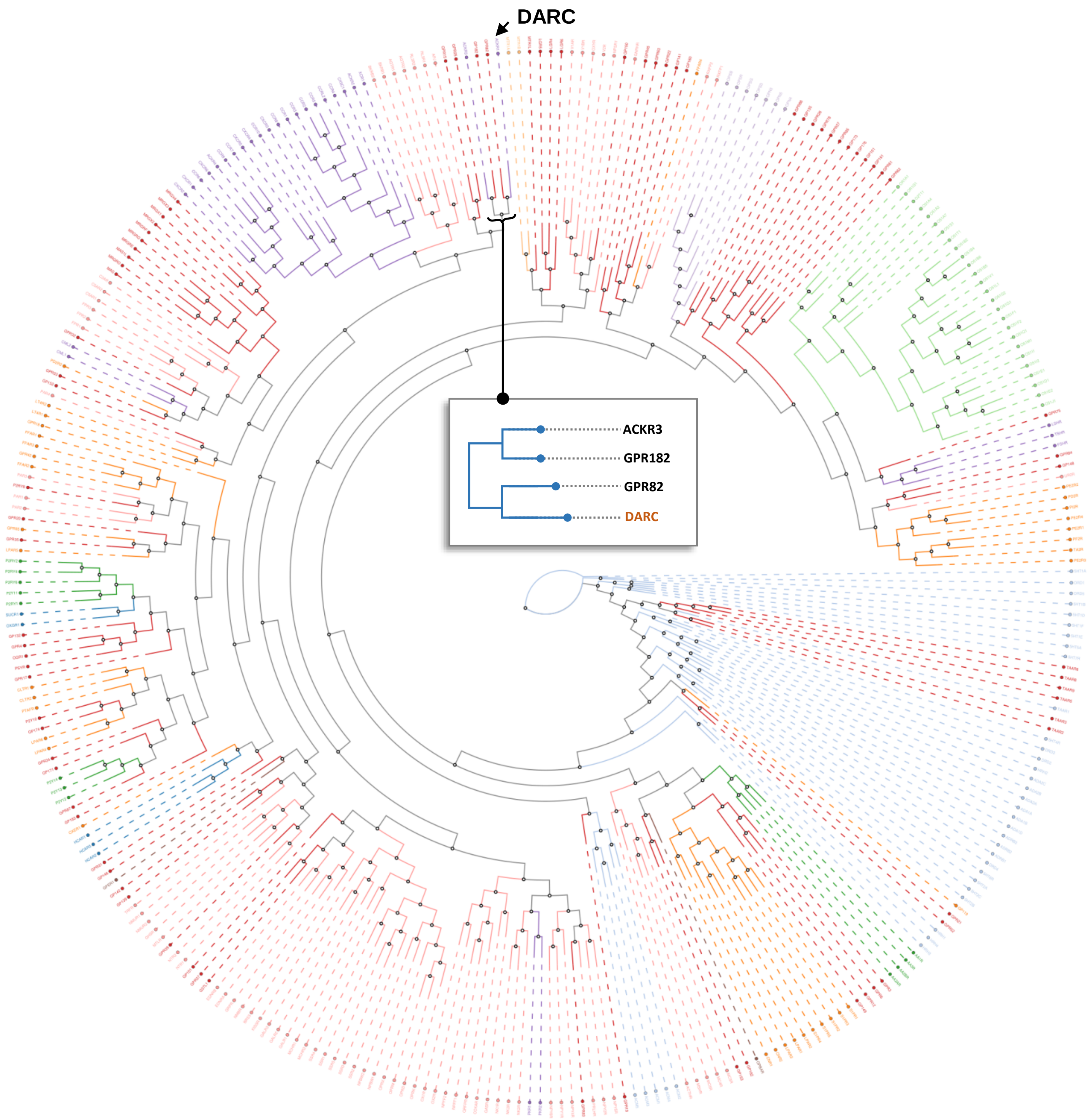


**Extended Data Fig. 1. Phylogenetic analysis of GPCRs highlighting the position of DARC.** (Phylogenetic plot obtained from www.gpcrdb.org)

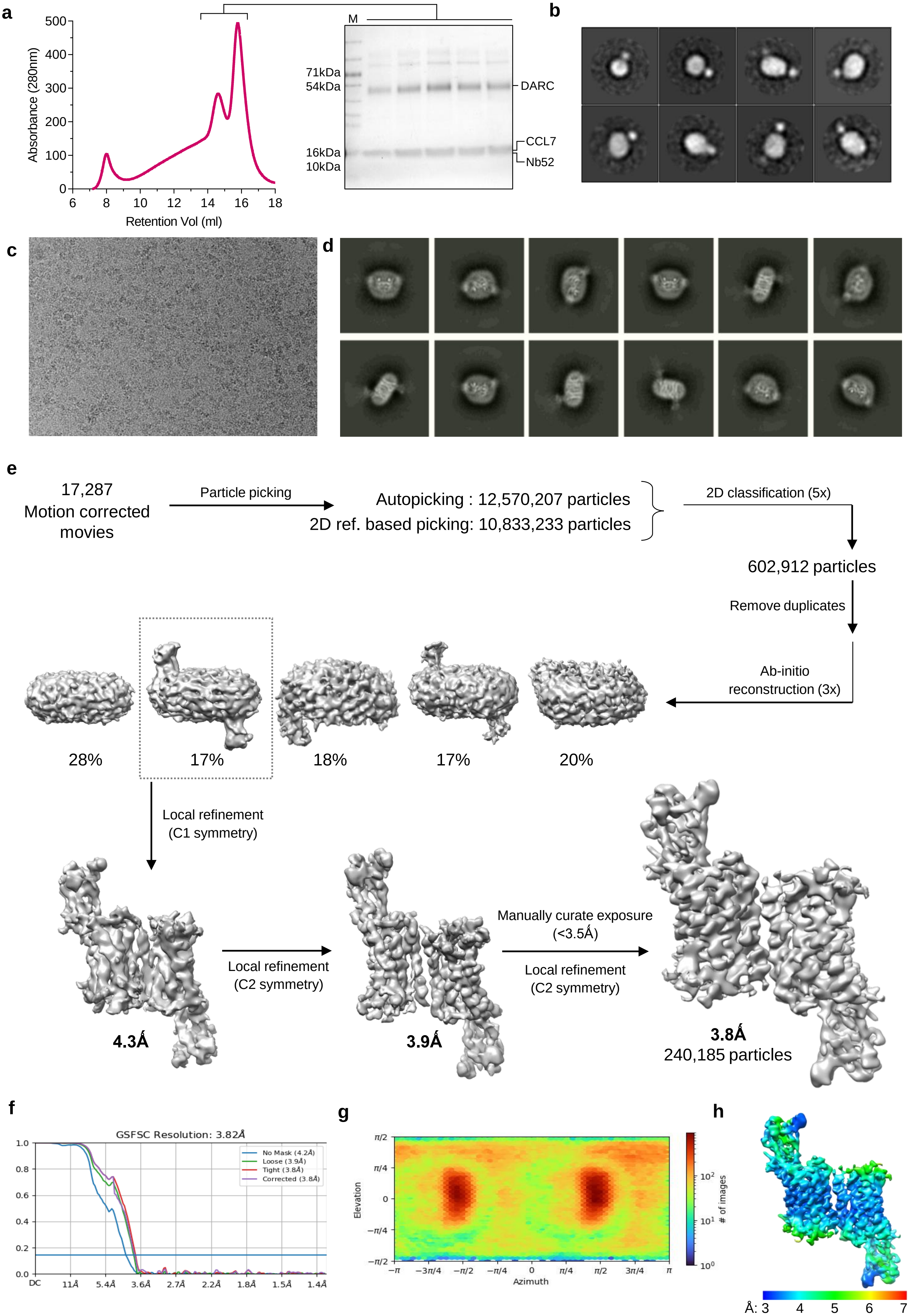
a
Absorbance (280nm)
Retention Vol (ml)
M
71kDa
54kDa
16kDa
10kDa
DARC
CCL7
Nb52
b
c
d
e
17,287
Motion corrected
movies
Particle picking
Autopicking : 12,570,207 particles
2D ref. based picking: 10,833,233 particles
2D classification (5x)
602,912 particles
Remove duplicates
Ab-initio
reconstruction (3x)
28%
17%
18%
17%
20%
Local refinement
(C1 symmetry)
4.3Å
Local refinement
(C2 symmetry)
3.9Å
Manually curate exposure
(<3.5Å)
Local refinement
(C2 symmetry)
3.8Å
240,185 particles
f
GSFSC Resolution: 3.82Å
No Mask (4.2Å)
Loose (3.9Å)
Tight (3.8Å)
Corrected (3.8Å)
g
Elevation
Azimuth
### of images
h
Å: 3 4 5 6 7

**Extended Data Fig. 2. Reconstitution and cryo-EM structure of the CCL7-DARC complex.**

**a.** Gel-filtration chromatogram and SDS-PAGE profile of Nb52-CCL7-DARC complex. **b.** Negative staining-EM 2D class averages. **c.** Representative motion corrected micrograph. **d** Selected 2D class averages. **e.** Schematic representation of cryo-EM data processing pipeline. **f.** Gold standard fourier shell correlation curve (GFSC) at 0.143 cut-off was used to determine the overall resolution of the map. **g.** Angular plot of the particles used for 3D reconstruction. **h.** Local resolution map of the 3D reconstruction in front view.

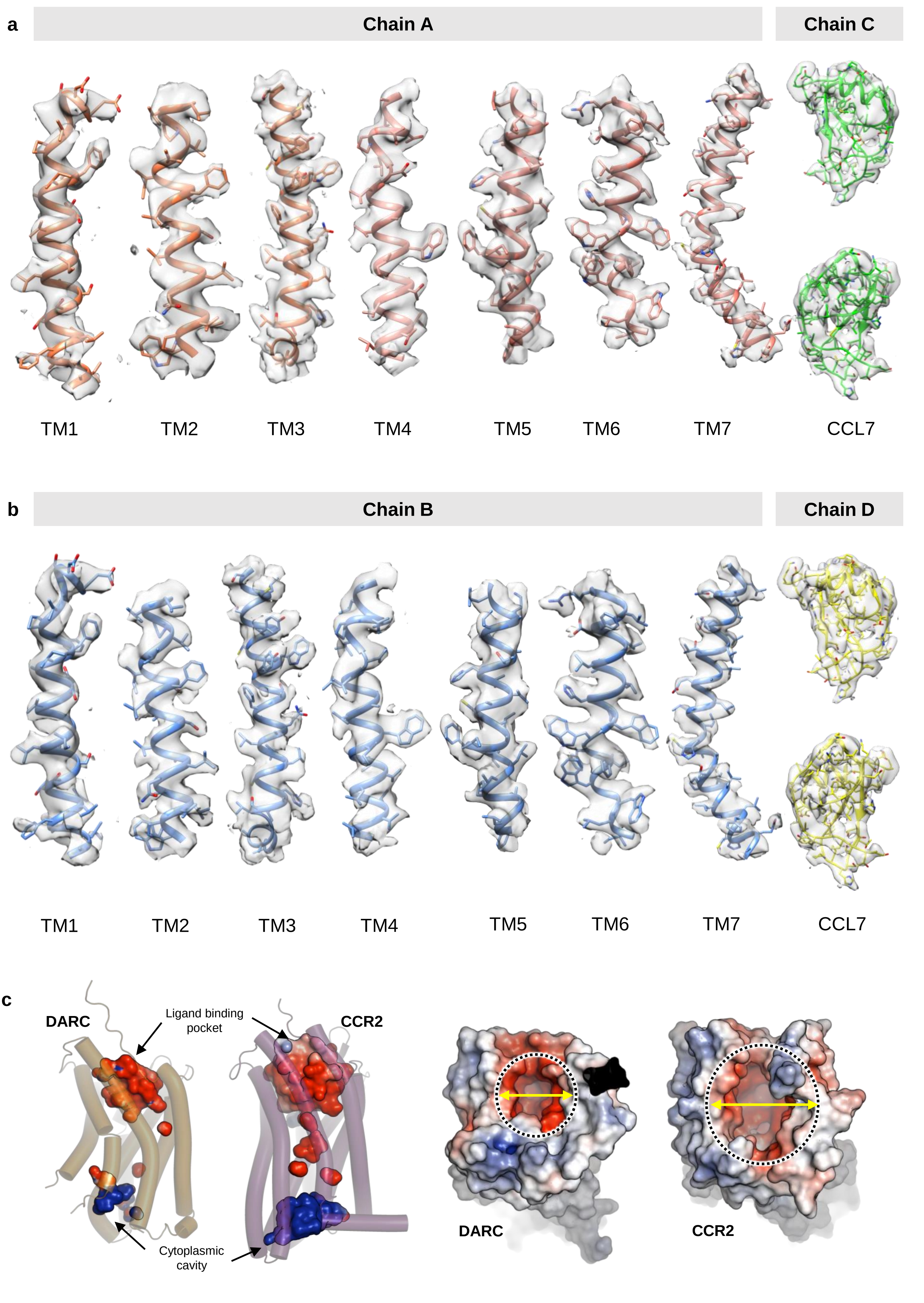
a
Chain A
Chain C
TM1
TM2
TM3
TM4
TM5
TM6
TM7
CCL7
b
Chain B
Chain D
TM1
TM2
TM3
TM4
TM5
TM6
TM7
CCL7
c
DARC
Ligand binding pocket
CCR2
Cytoplasmic cavity
DARC
CCR2

**Extended Data Fig. 3. Cryo-EM density maps and docking sites on DARC in comparison with CCR2.**

**a-b.** EM densities for TM helices and CCL7 in both the protomers are shown. **c.** The extracellular and intracellular cavities have been highlighted in CCL7 bound DARC in comparison with CCL2-CCR2 (PDB: 7XA3) (left). Receptors are shown as coulombic charged surface to depict the constricted ligand binding pocket on DARC (right).

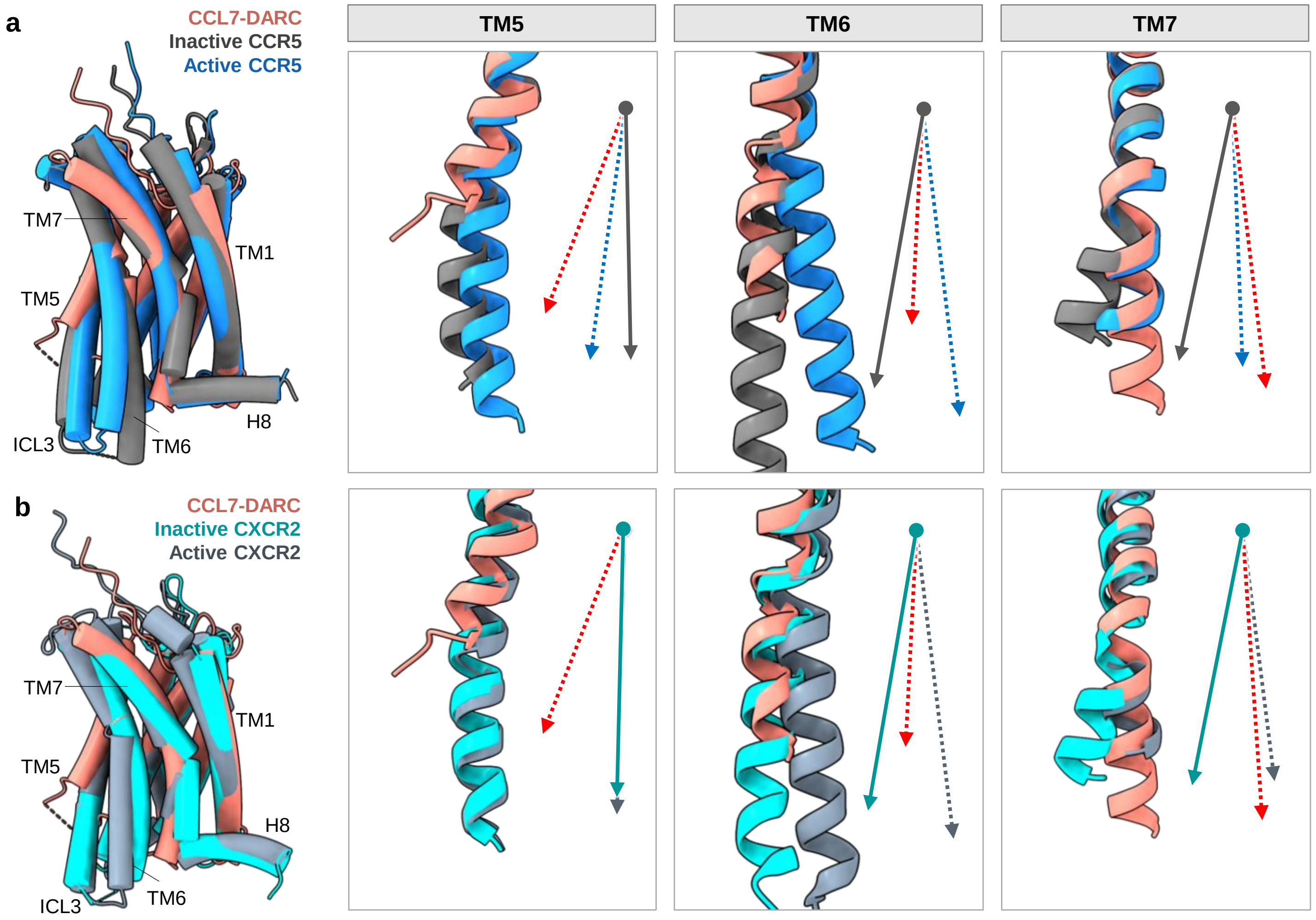


**Extended Data Fig. 4. Conformation of DARC in comparison with inactive and active state chemokine receptors.**

**a-b.** Structural alignment of CCL7-DARC with inactive CCR5 (PDB: 5UIW), CXCR2 (PDB: 6LFL) and active CCR5 (PDB: 7O7F), CXCR2 (PDB: 6LFO) structures (left). The deviations in TM5, TM6 and TM7 are shown. (Arrows with gray and cyan depict inactive CCR5 and CXCR2, respectively. Arrows with blue and slate gray depict active CCR5 and CXCR2, respectively. TMs of DARC have been shown in red arrow).

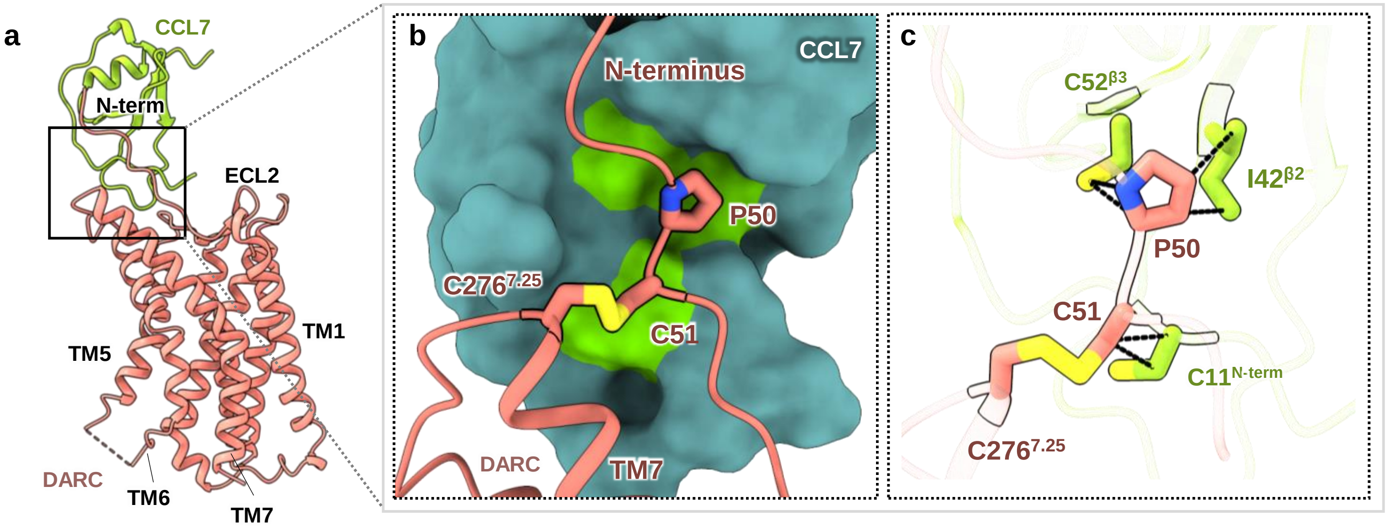


**Extended Data Fig. 5. The interface of P-C motif on DARC with chemokines.**

**a.** CCL7 bound to DARC is shown as ribbon representation. **b.** Residues of the P-C motif are shown as sticks bound to CCL7 as surface. The P-C motif residues dock into the cavity in CCL7 formed by the C-I-C residues. **c.** Non-bonded contacts between P-C and C-I-C residues are shown as black dashed lines.

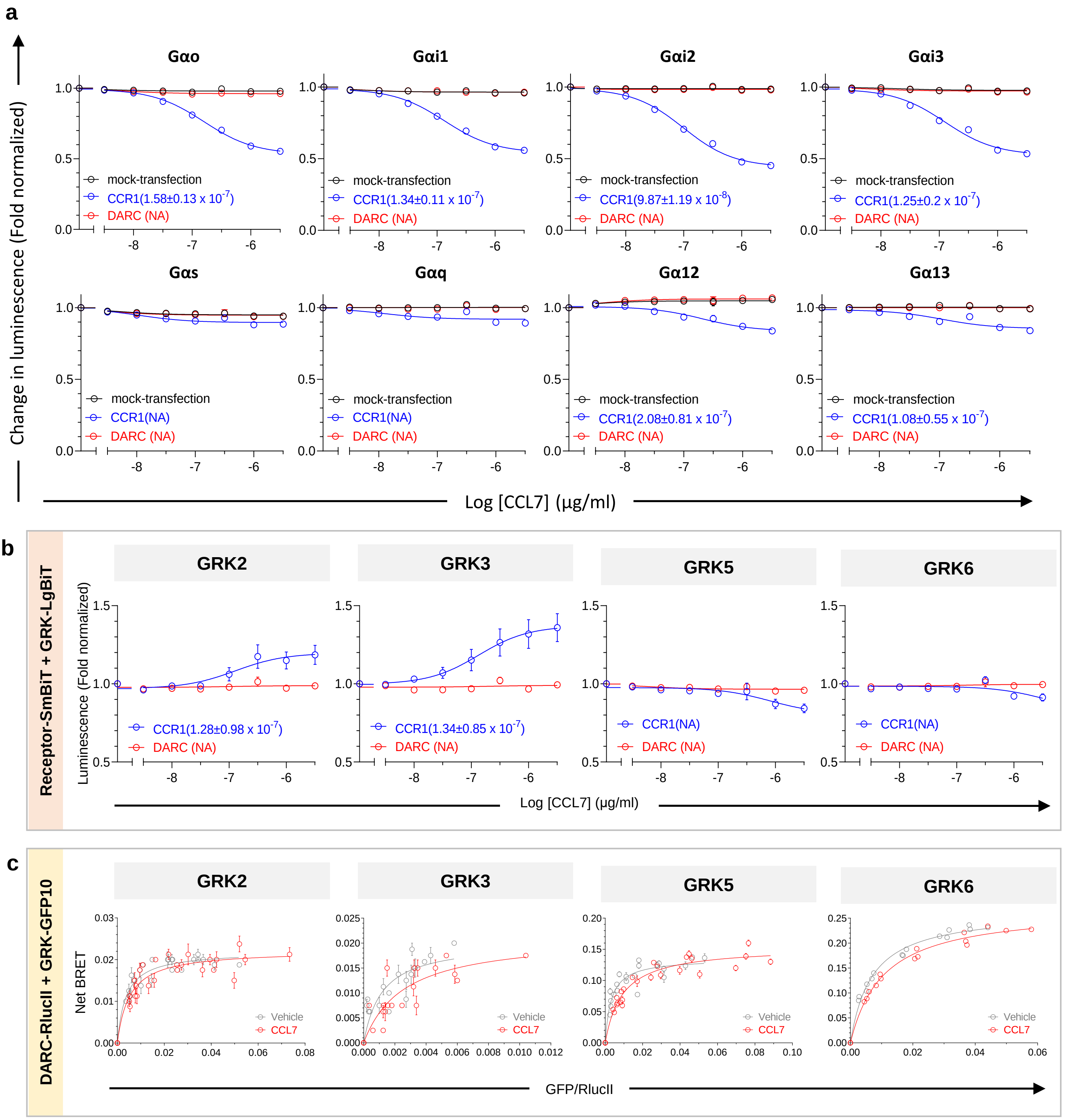


**Extended Data Fig. 6. Lack of G protein activation and GRK recruitment downstream to DARC.**

**a.** Stimulation with CCL7 does not induce G protein heterotrimer dissociation. Data (mean±SEM) represent three independent experiments, normalized with respect to baseline signal (i.e., vehicle treatment) for each set. **b-c.** CCL7 treatment fails to induce GRK recruitment to DARC, as measured by NanoBiT assay (a) and BRET assay (b). Data (mean±SEM) represent three independent experiments, normalized with respect to baseline signal (i.e., vehicle treatment) for each set, treated as 1 (a). Data (mean±SEM) represent four to six independent experiments (b).

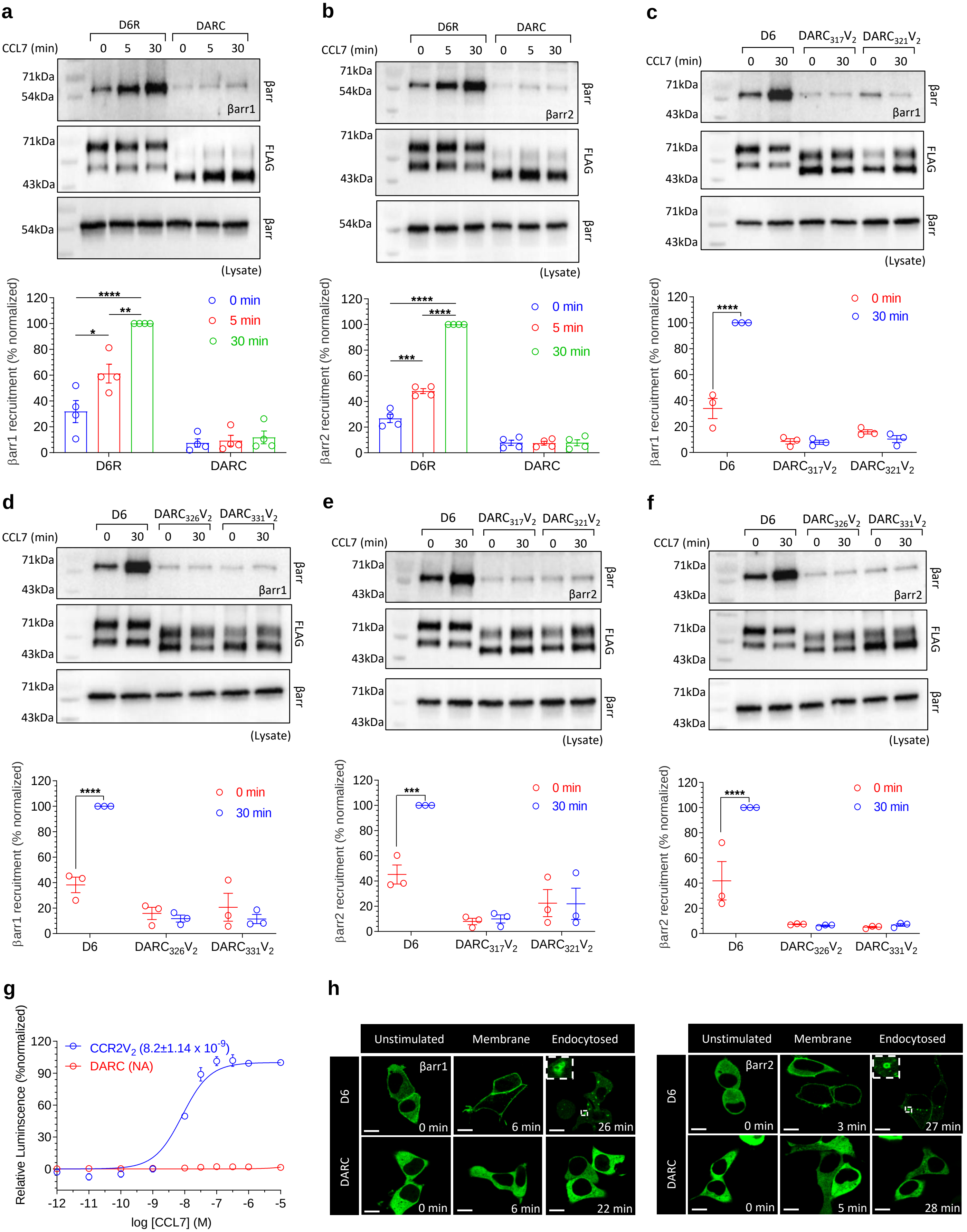

a
D6R
DARC
CCL7 (min)
0 5 30 0 5 30
71kDa
54kDa
βarr
βarr1
FLAG
43kDa
(Lysate)
βarr1 recruitment (% normalized)
0 min
5 min
30 min
D6R
DARC
b
βarr2
βarr2 recruitment (% normalized)
c
D6
DARC317V2
DARC321V2
d
DARC326V2
DARC331V2
e
f
g
CCR2V2 (8.2±1.14 x 10-9)
DARC (NA)
Relative Luminscence (%normalized)
log [CCL7] (M)
h
Unstimulated
Membrane
Endocytosed
D6
DARC
0 min
6 min
26 min
22 min
3 min
27 min
5 min
28 min

**Extended Data Fig. 7. Lack of β-arrestin recruitment to DARC.**

**a-b.** DARC fails to elicit β-arrestin recruitment in response to CCL7, as measured by co-immunoprecipitation. A representative image from four independent experiments and densitometry-based quantification of data (mean±SEM) normalized with respect to the signal observed at 30mins after ligand stimulation of D6R (treated as 100%) is shown here. Data are analysed using two-way ANOVA (Sidak's multiple comparison; *p < 0.05, **p < 0.01, ***p < 0.001, ****p < 0.0001). **c-f.** Addition of $V_2$-tail at different positions fails to induce β-arrestin recruitment. A representative image from four independent experiments and densitometry-based quantification of data (mean±SEM) normalized with respect to the signal observed at 30mins after ligand stimulation of D6R (treated as 100%) is shown here. Data are analysed using two-way ANOVA (Sidak's multiple comparison; ***p < 0.001, ****p < 0.0001). **g.** TANGO assay confirms the same. Data (mean±SEM) represents four independent experiments normalized with respect to the highest signal observed for CCR2$V_2$ (treated as 100%). **h.** Confocal assay corroborates the lack of β-arrestin recruitment and trafficking downstream to DARC. A representative image visualizing mYFP-β-arrestin is shown from two independent experiments (scale bar 10μm).

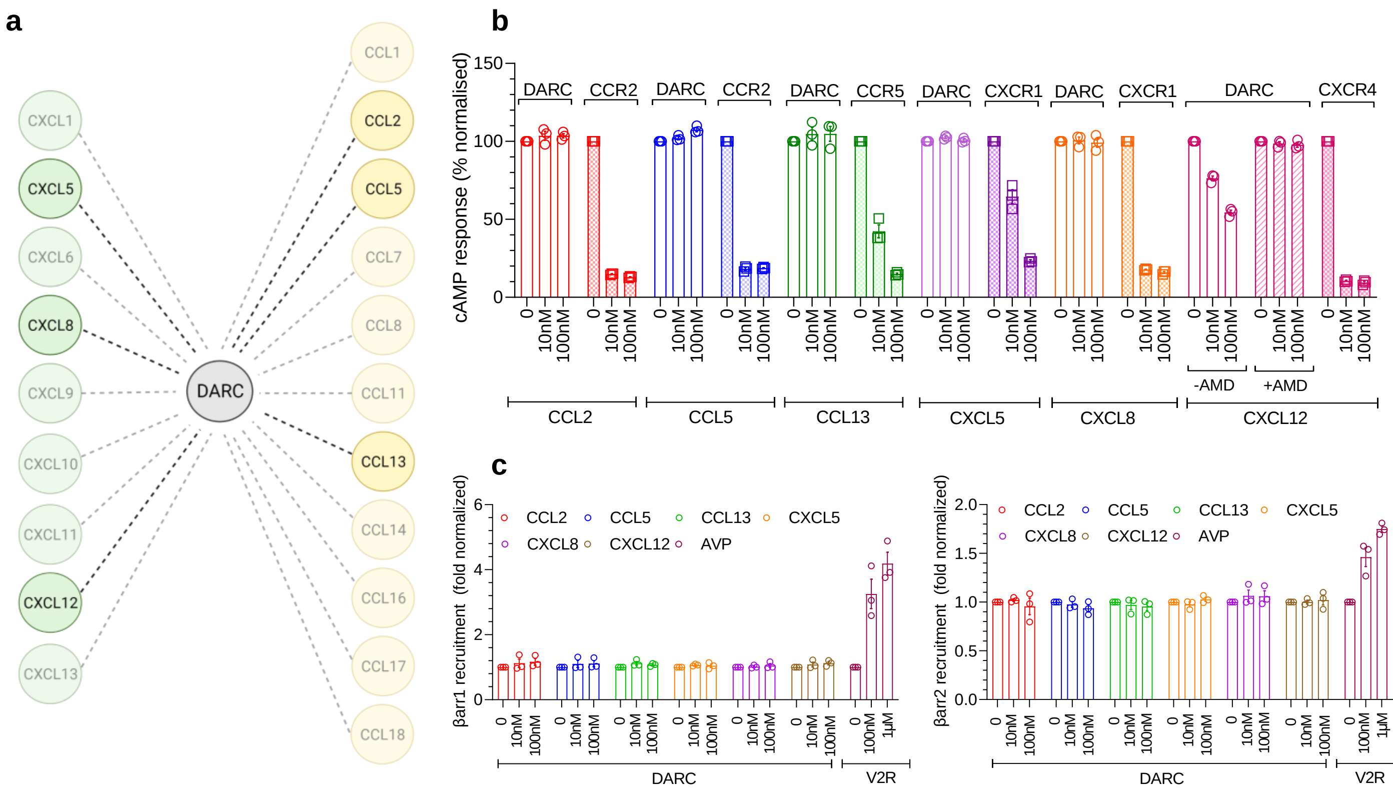


**Extended Data Fig. 8. Multiple chemokines fail to activate G protein signaling and β-arrestin recruitment downstream to DARC.**

**a.** Schematic representation of the different chemokines that bind DARC (chemokines used in subsequent assays are highlighted). **b.** No decrease in cytosolic cAMP is observed upon stimulating DARC with multiple CC and CXC chemokines. Data (mean±SEM) represents three independent experiments, normalized with respect to the baseline signal observed for each set (treated as 100%). AMD refers to AMD3100, a selective CXCR4 antagonist, and it is used here to block CXCL12-induced cAMP response arising from the endogenous CXCR4 in HEK2-93 cells. **c.** Multiple CC and CXC chemokines tested fail to induce β-arrestin recruitment to DARC, as measured by NanoBiT assay. Data (mean±SEM) represents three independent experiments, normalized with respect to the baseline signal observed for each set (treated as 1).

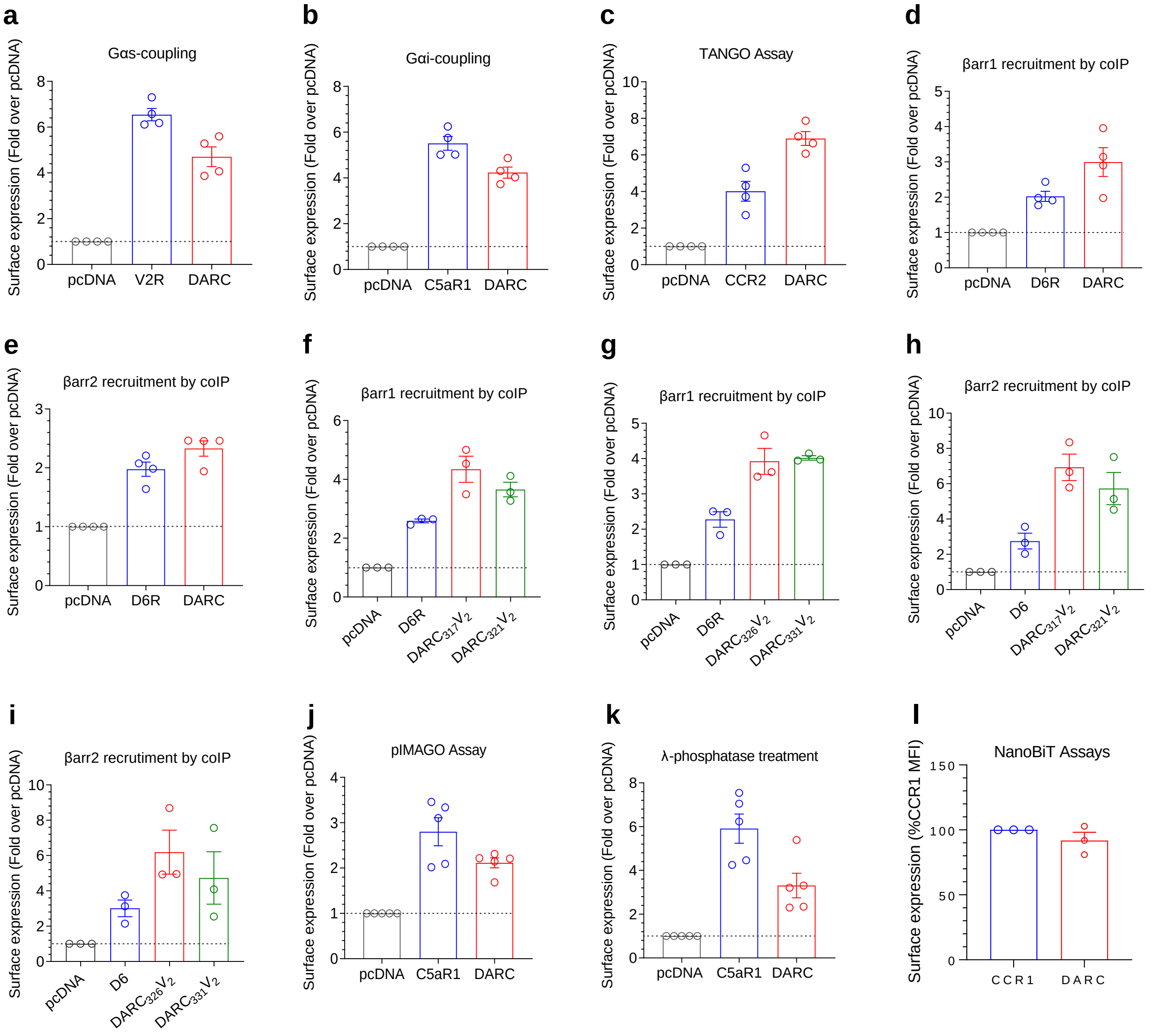


**Extended Data Fig. 9. Surface expression of receptors.**

**a-l.** Surface expression of all receptors as measured by whole-cell ELISA for various assays.
